# Stress from environmental change drives clearance of a persistent enteric virus

**DOI:** 10.1101/2024.11.06.622373

**Authors:** Christin Herrmann, Kimberly Zaldana, Eva L Agostino, Sergei B Koralov, Ken Cadwell

## Abstract

Persistent viral infections are associated with long-term health issues and prolonged transmission. How external perturbations after initial exposure affect the duration of infection is unclear. We discovered that murine astrovirus, an enteric RNA virus, persists indefinitely when mice remain unperturbed but is cleared rapidly after cage change. Besides eliminating the external viral reservoir, cage change also induced a transcriptional defense response in the intestinal epithelium. We further identified that displacing infected animals initially caused a temporary period of immune suppression through the stress hormone corticosterone, which was followed by an immune rebound characterized by an increase in CD8 T cells responsible for the epithelial antiviral responses. Our findings show how viral persistence can be disrupted by preventing re-exposure and activating immunity upon stress recovery, indicating that external factors can be manipulated to shorten the duration of a viral infection.

## Introduction

The time course of infection for many viruses, such as RNA viruses that cause self-limiting respiratory or gastrointestinal diseases, is characterized by a brief period of peak viral burden followed by resolution within days. However, the development of more sensitive techniques have enabled detection of viral genomes, replication intermediates, and protein antigens weeks and even months after the initial infection for a growing list of viruses previously considered non-persistent^1-13^. Prolonged infection may contribute to long-term complications, such as post-acute sequalae of SARS-CoV-2 (long COVID) or increased opportunities for the virus to accumulate mutations and transmit to another host^13-18^.

Evasion of CD8 T cell responses is a hallmark of chronic infections^19^. However, unlike retroviruses and certain DNA viruses that display periods of latency, non-integrating RNA viruses typically exhibit continuous replication and production of MHC antigens. During persistent lymphocytic choriomeningitis virus (LCMV) infection of mice, unremitting stimulation of CD8 T cells induces inhibitory receptors such as PD-1, leading to a state of exhaustion characterized by reduced activity^20,21^. T cells specific for hepatitis B virus (HBV) and HCV in humans display similar signs of exhaustion^19,22,23^. In the gut, CD8 T cells fail to recognize immune-privileged tuft cells that are infected by persistent strains of murine norovirus (MNV)^24-26^. Therapeutic blockade of PD-1 and innate immune activation through IFNλ administration overcome these failures to clear LCMV and MNV, respectively, paving the way for similar strategies in humans^27,28^. Thus, identifying determinants of persistence upstream of CD8 T cells may reveal opportunities for non-pharmaceutical interventions, which would be more facile to implement as a public health measure during an outbreak.

Astroviruses are non-enveloped positive strand RNA viruses that infect a wide range of animal species^29^. In humans, astroviruses are a common cause of an acute, self-limiting gastroenteritis, and more than 91% of children have experienced at least one infection by age five^30^. Astroviruses were discovered in lab and wild mice in 2012^31,32^, and subsequently shown to spread fecal-orally and infect goblet cells in the intestine similar to human astroviruses^33,34^. Murine astrovirus-1 (MuAstV) is endemic in many mouse facilities where it persists asymptomatically in immunocompromised mice^32,35,36^. In contrast, it has been reported that the virus is cleared within a couple weeks in immunocompetent wild-type (WT) mice^35,36^. In this study, we found that MuAstV is only cleared because of animal facility cage change procedures and can persistently infect WT mice if they remain in the same cage for the duration of the experiment. We show that cage change mediates MuAstV clearance by preventing reinfection while simultaneously inducing a short inflammatory burst through a stress-induced antiviral response by CD8 T cells and the intestinal epithelium. These findings demonstrate that environmental conditions determine the duration of a viral infection.

## Results

### MuAstV establishes a persistent infection in the absence of cage change

Germ-free (GF) mice require less frequent exchange of cages to remove dirty bedding. While investigating the role of the microbiota during MuAstV infection in WT mice, we observed that the timing of virus decline in infected GF mice was variable between experiments and coincided with cage change. To test whether cage change was responsible for MuAstV clearance, we compared viral shedding in the stool over time in WT mice with and without cage change at 14 days post inoculation (dpi). Viral RNA (vRNA) was below the limit of detection by day 21 in mice that experienced cage change, while mice without a cage change shed virus in their stool throughout the experiment (nearly 2 months) (**Figure 1A**). Similar to GF conditions, specific pathogen free (SPF) mice also cleared the virus by 28dpi following cage change at 14dpi and remained infected when maintained in the same cage (**Figure 1B, Figure S1A**). When we investigated the timing of MuAstV decrease through daily monitoring after cage change, we found that virus levels were reduced more rapidly in SPF mice and that GF mice also cleared the virus by 3dpi (**Figure 1C-D**). Thus, cage change mediates viral clearance in both the presence and absence of the microbiota, and we decided to use both GF and SPF mice to investigate mechanism (as indicated in figure legends).

**Figure 1.**
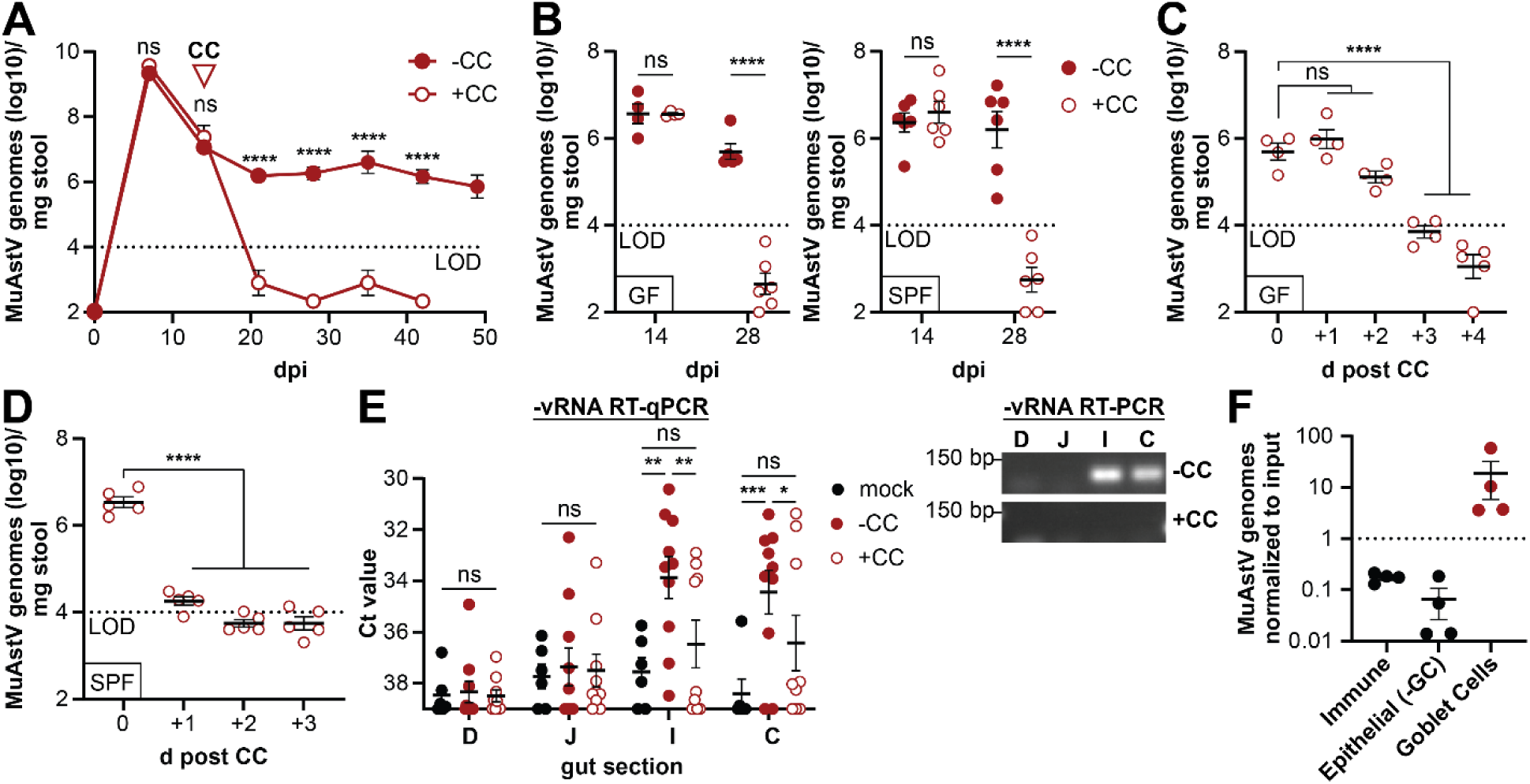
MuAstV establishes a persistent infection in the absence of cage change: **(A)** MuAstV genome levels in stool of germ-free (GF) B6 mice over time +/- cage change (CC) at 14 days post inoculation (dpi) detected by RT-qPCR. Mice were inoculated with 1E10 viral genomes via oral gavage. n= 3 mice. **(B)** MuAstV genome levels in GF and specific-pathogen-free (SPF) mice +/-CC. n = 5-6 mice. **(C-D)** MuAstV genome levels in stool from GF (C) and SPF (D) mice measured daily after CC. n = 4-5 mice. **(E)** Ct values for MuAstV negative strand (-vRNA) RT-qPCR and matching DNA gel of negative strand RT-PCR in different gut sections of mock controls and MuAstV infected mice +/-CC. D - Duodenum, J - Jejunum, I - Ileum, C - Colon. Data pooled from GF and SPF mice. n = 4-10 mice. **(F)** MuAstV genome levels by RT-qPCR detected in cell populations sorted from the ileum at 14dpi of SPF mice (no CC). Immune: CD45+, Epithelial (- GC): EpCAM+CLCA1-, Goblet Cells: EpCAM+CLCA1+. n = 4 mice. Each dot represents data from one animal in panels (B-F). All data are represented as mean ± SEM from at least two independent experiments. LOD = limit of detection. ns = not significant, *p≤0.05, **p≤0.01, ****p≤0.0001 by 2way ANOVA and Holm-Šídák’s comparison for (A)-(E). See also Figure S1.

It is unlikely that the prolonged presence of viral genomes in stool represents continuous uptake of the same non-replicating virus from consumption of stool (coprophagia) because we would expect attrition over time. As plaque assays are not available for this virus, we analyzed gut tissue for presence of negative strand vRNA (–vRNA), a replication intermediate of positive strand RNA viruses^29^. As a positive control, we analyzed tissue at 7dpi and found that –vRNA was readily detected throughout the small intestine and colon well above background levels detected in mock infected mice (**Figure S1B**). At 21dpi, –vRNA were still detectable in the ileum and colon of mice without cage change, while cage change at 14dpi resulted in levels comparable to mock infected mice (**Figure 1E**). Only the ileum and colon sections of mice without cage change had visible bands for –vRNA when we analyzed RT-PCR products (**Figure 1E**). Consistent with studies showing that MuAstV replicates in goblet cells at peak of infection^33,34^, we found that MuAstV genomes were enriched in sorted goblet cells (EpCAM+CLCA1+) compared to the immune cell fraction (CD45+) or epithelial fraction without goblet cells (EpCAM+CLCA1-) (**Figure 1F and S1C**). These results indicate that MuAstV persistence represents ongoing viral replication, and that goblet cells remain a major target cell at a timepoint that is sensitive to cage change-mediated clearance.

### Continued exposure contributes to persistent infection

To determine whether the input quantity of virus affects persistence, we inoculated mice with a dilution series of MuAstV and quantified viral shedding for 4 weeks without cage change. Inoculation with concentrations ranging from 1E7 to 1E10 viral genome copies led to similar levels of virus in stool with high levels detected at 7dpi followed by persistence. Administering 1E6 genomes did not lead to successful infection in all mice, but we observed continuous shedding among those that were positive for vRNA at earlier time points (**Figure 2A, Figure S1D**). These results demonstrate that, in the absence of cage change, mice remain persistently infected as long as the initial infection occurs.

**Figure 2.**
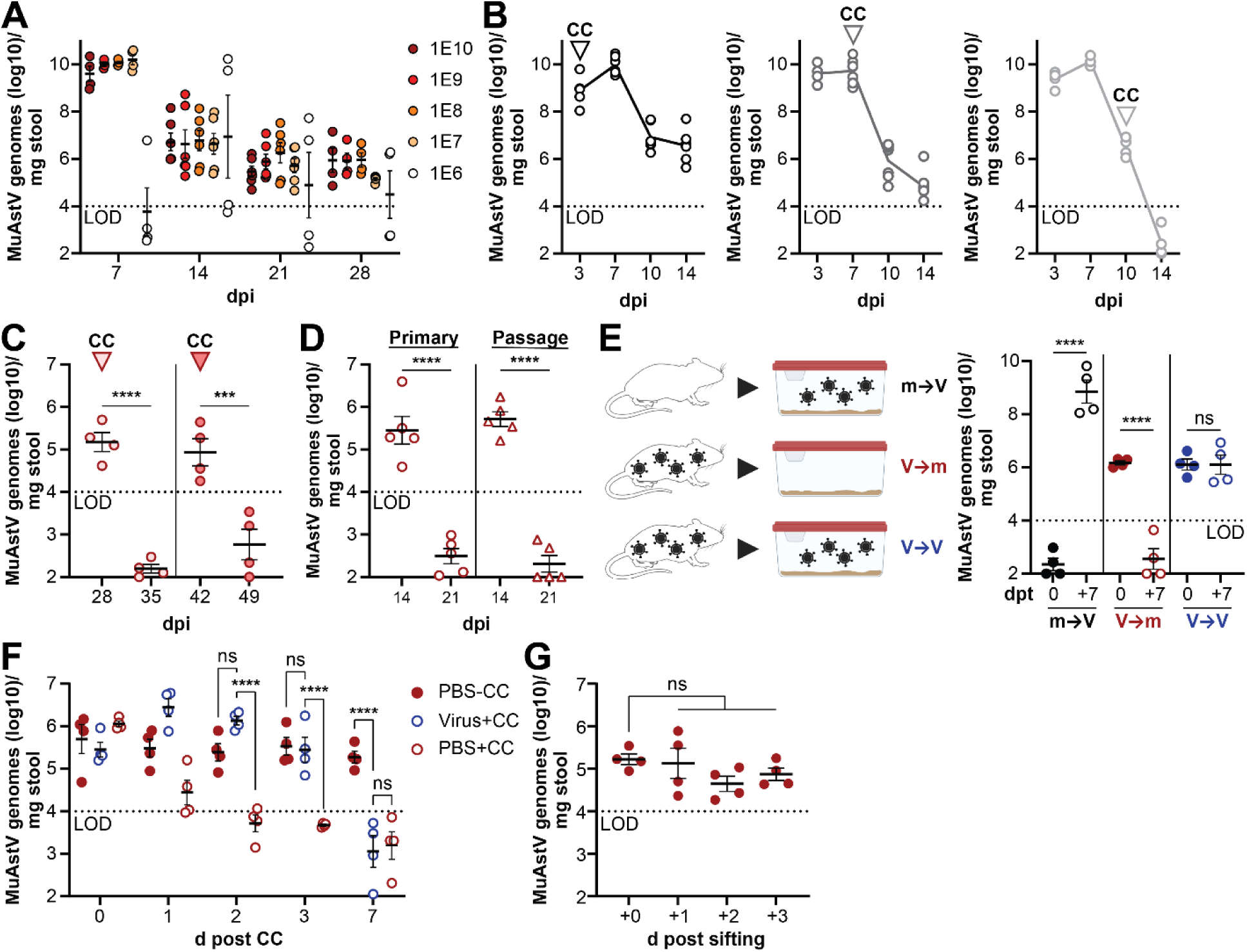
Continued exposure contributes to persistent infection: **(A)** Time course of MuAstV levels in stool detected by RT-qPCR from mice inoculated with 10-fold dilutions of the viral inoculum (1E6 – 1E10 genome copies) in the absence of CC. n = 4-6 mice. **(B)** MuAstV levels over time after CC on 3, 7 or 10dpi. n = 4-7 mice. **(C)** MuAstV levels before and after CC at either 28 or 42dpi. n = 4 mice. **(D)** MuAstV levels before and after CC in primary infection or after passage of the virus. n = 4 mice. **(E)** MuAstV levels before and after cage ‘trade’ at 14dpi. Groups: m➔V = mock mice transferred into cages previously containing virus-infected mice, V➔m = infected mice into cages previously containing mock mice. V➔V = infected mice into cages previously containing a separate cohort of virus-infected mice. dpt = day post trade. n = 4 mice **(F)** MuAstV levels in mice with repeated oral gavage of virus or PBS at 0h, 6h, 12h and 24h after cage change. n = 4 mice. **(G)** MuAstV levels in mice before and after sifting the bedding. N = 4 mice. Data from GF and SPF mice were pooled in panels (A) and (E). Only SPF mice were used for all others. Each dot represents data from one animal in all panels. All data are represented as mean ± SEM from at least two independent experiments. ns = not significant, ***p≤0.001, ****p≤0.0001 by 2way ANOVA and Holm-Šídák’s comparison. See also Figure S1.

Next, we investigated the impact of the timing of cage change. Cage change at 3dpi and 7dpi did not eliminate viral persistence. However, cage change at 10dpi led to rapid viral clearance (**Figure 2B**). Although it is possible that the virus acquires mutations over time that facilitate persistence, changing cages later at 28 or 42dpi still resulted in reduction of vRNA levels below the limit of detection (**Figure 2C**). To further demonstrate that cage change and not mutation of the virus is the determining factor of persistence, we co-housed infected mice at 14dpi with naïve mice for another 14 days and then performed the cage change on the second set of mice. Passaging the virus through mice did not alter viral clearance (**Figure 2D**). This indicates that even if given time to evolve, MuAstV remains susceptible to clearance following cage change. Subsequent experiments were performed with 14dpi as standard time of cage change.

We tested whether continued exposure to the virus in an unchanged cage contributes to persistence by swapping cages between infected and mock infected mice (**Figure 2E**). As expected, mock infected mice placed into cages that previously harbored virus-infected mice (**m**➔**V**) became MuAstV-positive, indicating that infectious virus was retained in the cage after removal of the infected mice. Infected mice placed into cages previously containing mock-infected mice (**V**➔**m**) cleared the virus similar to a cage change. Infected mice placed into cages previously containing virus-infected mice (**V**➔**V**) remained MuAstV-positive, indicating that presence of virus in the environment contributes to persistence.

To assess whether continued exposure to virus is sufficient to mediate persistence, we repeatedly gavaged mice at 0, 6, 12, and 24h post cage change with either phosphate-buffered saline (PBS) or virus after cage change (**Figure 2F**). PBS-treated mice without cage change had unaltered virus levels. Repetitive administration of MuAstV delayed but did not prevent clearance of virus compared with PBS treatment following cage change, suggesting an additional component of changing cage, such as stress caused by the new environment, affects persistence^37^. Consistent with this possibility, mice kept in the same cage remained infected after we removed stool pellets by sifting the bedding (**Figure 2G and S1E**). Together, these data indicate that continued exposure contributes to MuAstV persistence, but that other factors associated with changing cages are necessary for viral clearance.

### CD8 T cells are necessary for cage change-mediated MuAstV clearance

The timing at which the cage change is effective in mediating clearance (after day 7) coincides with the expected start of adaptive immune responses (**Figure 2B**). Mice that are infected and clear MuAstV display protection from immediate reinfection^35^. We verified the development of protective immunity in our system by reinfecting mice at 28dpi. Infected mice without cage change remained persistently infected after a second inoculation, while mice that underwent cage change did not become reinfected after clearing the initial infection (**Figure 3A**). We examined humoral responses to the immunodominant spike domain by developing an ELISA to measure antibodies specific to our MuAstV strain based on a 3D structure predicted by AlphaFold^38-41^ (**Figure S2A**). Although MuAstV infected mice had high levels of virus-specific IgA in stool and IgG in serum at 28dpi compared with mock infected mice, these antibody levels were not affected by cage change (**Figure 3B and S2B**). Virus-specific antibodies were detectable before cage change, sometime between 7 and 14dpi (**Figure 3B**). Additionally, we did not find evidence that cage change enhanced the development of neutralizing antibodies as viral inoculum incubated with serum from mock infected mice, MuAstV infected mice, and MuAstV infected mice with cage change were all still infectious (**Figure 3C**). Together, these experiments do not support an effect of cage change on antiviral antibody activity.

**Figure 3.**
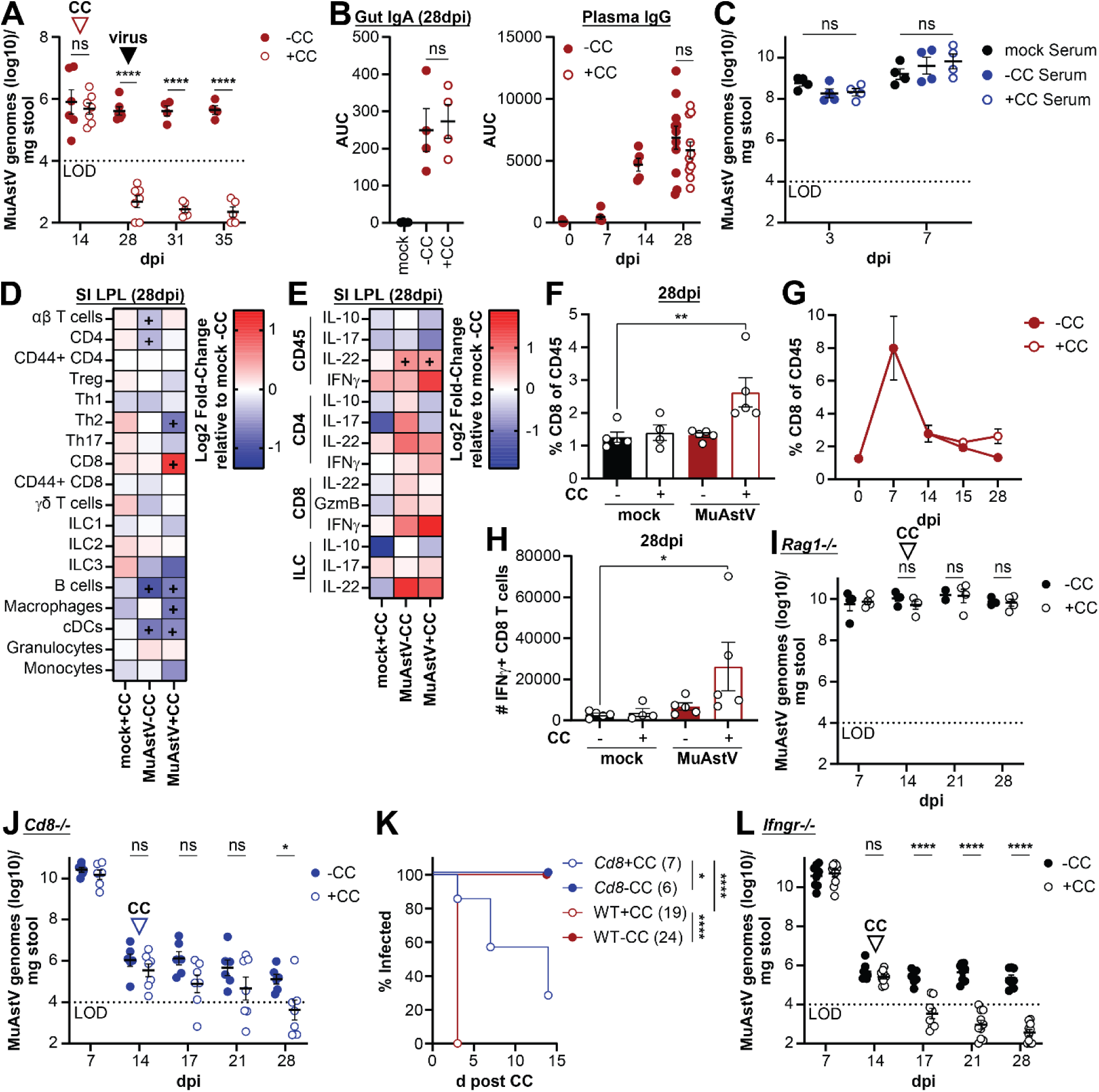
CD8 T cells are necessary for cage change-mediated MuAstV clearance: **(A)** MuAstV genome levels over time +/-CC at 14dpi and reinfection with the same dose of MuAstV 28dpi (virus). n = 4-7 mice. **(B)** Area under the curve (AUC) for gut anti-spike IgA levels in the ileum and time course of plasma anti-spike IgG from mice with MuAstV +/-CC at 14dpi measured by ELISA. n = 4-11 mice. **(C)** MuAstV levels in stool of mice infected with virus inoculum incubated with serum collected at 28dpi from mock treated mice or MuAstV infected mice +/-CC. n = 4 mice. **(D-E)** Heatmaps of average fold changes in indicated groups compared to mock-CC for immune population frequencies (D) and frequency of cytokine production (E) in the lamina propria leukocytes (LPL) of the small intestine (SI) of GF mice as measured by flow cytometry. n = 4-5 mice. **(F)** Percentage of CD8 T cells of CD45+ cells in SI LPL. n = 4-5 mice. **(G)** Time course for the percentage of CD8 T cells out of CD45+ cells in the SI LPL of infected mice +/-CC at 14dpi. N = 2-5 mice. **(H)** Number of IFNγ+ CD8 T cells in SI LPL. n = 4-5 mice. **(I-J)** MuAstV levels over time in *Rag1^-/-^* (I) or *Cd8^-/-^* (J) mice +/- CC at 14dpi. n = 4-7 mice. **(K)** Comparison of % of infected mice remaining in WT and *Cd8^-/-^* mice +/-CC. **(L)** MuAstV levels over time in *Ifngr^-/-^* mice +/-CC at 14dpi. n = 8-10 mice. Data from GF and SPF mice were pooled in panels (A-C), only GF mice were for panels (D-H) and only SPF mice for panels (I-L). Each dot represents data from one animal in panels (A-C), (F-J) and (L). All data are represented as mean ± SEM from at least two independent experiments. ns = not significant, *p≤0.05, **p≤0.01, ****p≤0.0001 by 2way ANOVA and Holm-Šídák’s comparison for (A-C), (F-J) and (L), two-sided Student’s T-test for (D+E), and Mantel-Cox analysis for (K). For panel (D+E) ‘+’ denotes a fold change with any p≤0.05. See also Figure S2.

To more broadly examine how the immune system responds to cage change during MuAstV infection, we analyzed lamina propria leukocytes (LPL) from the small intestine and colon by flow cytometry. We used GF mice for this analysis because overlapping responses to other microbes might interfere with our ability to identify reactions that are most relevant to the virus^42^. When compared with mock infected mice, we observed alterations in the proportion of several immune cell populations following MuAstV infection at multiple time points in both organs in the absence of cage change. This included a reduction in B cells and upregulation of IL-22 producing cells (**Figure 3D-E and S3C-D**), as previously reported^42^. Infected mice with cage change was largely similar, except displayed an additional increase in CD8 T cells and IFNγ+ CD8 T cells at 28dpi (**Figure 3D-H and S2E-F**). When we plotted the CD8 T cell levels over time, we observed that MuAstV infection led to a large increase at 7dpi that returned to baseline level in the no cage change group but remained above pre-infection levels in mice with cage change (**Figure 3G and S2C**).

We analyzed immunocompromised mice to further examine the contribution of adaptive immunity to cage change. We found that many of these mouse lines, either purchased from a vendor or bred in our institutional facility, were already infected with MuAstV. Therefore, we established virus-naïve lines through a combination of GF-rederivation, early cross fostering, or breeding through WT intermediates (see methods). Consistent with previous reports showing that *Rag1^-/-^* mice lacking mature T and B cells are persistently infected with MuAstV^32^, we found that virus levels were unresponsive to cage change when we infected naïve *Rag1^-/-^* mice (**Figure 3I**). However, we note that viral levels remained high in *Rag1^-/-^* mice after the typical peak on 7dpi, and thus, we cannot distinguish a role for lymphocytes that precedes cage change from a potential role in mediating clearance after cage change. As a less drastic form of compromised immunity, we tested mice deficient in CD8 T cells and IFNγ. In the absence of cage change, MuAstV levels peaked at 7dpi and then decreased in *Cd8^-/-^* mice to reach stable persistence, similar to WT mice (**Figure 3J**). Strikingly, MuAstV clearance was substantially delayed or failed to occur altogether in most *Cd8^-/-^* mice after cage change (**Figure 3J-K**). In contrast, mice lacking the IFNγ receptor effectively cleared MuAstV after cage change (**Figure 3L**). Similarly, blocking IFNγ with neutralizing antibodies did not alter the time course by which cage change mediated decreases in virus in both SPF and GF mice (**Figure S2G**). These results implicate the increase in gut CD8 T cells, and potentially other components of the adaptive immune system, in mediating MuAstV clearance induced by cage change. The increase in IFNγ production may be a marker of enhanced effector capacity but is dispensable or redundant for CD8 T cell-mediated control of MuAstV infection after cage change.

### Virus-specific CD8 T cells are not exhausted despite persistent infection

To identify MuAstV-specific CD8 T cells, we generated an MHC-I tetramer. First, we created a peptide library containing the 50 top predicted MHC peptides. This peptide pool stimulated IFNγ production in CD8 T cells in the small intestine LPL fraction in MuAstV infected mice but not in mock controls (**Figure 4A and S3A**). We then subdivided the library into pools of 5-16 peptides and found that only one reproduced this stimulatory activity. Testing the individual peptides represented in this pool identified two adjacent peptides (**Figure S3B-D**). The overlap between these peptides contains a 9 amino acid motif (AAIWNPIVV) predicted to bind MHC class I allele H-2-Db. This peptide alone stimulated IFNγ production by CD8 T cells similarly to the full peptide pool (**Figure 4A**).

**Figure 4.**
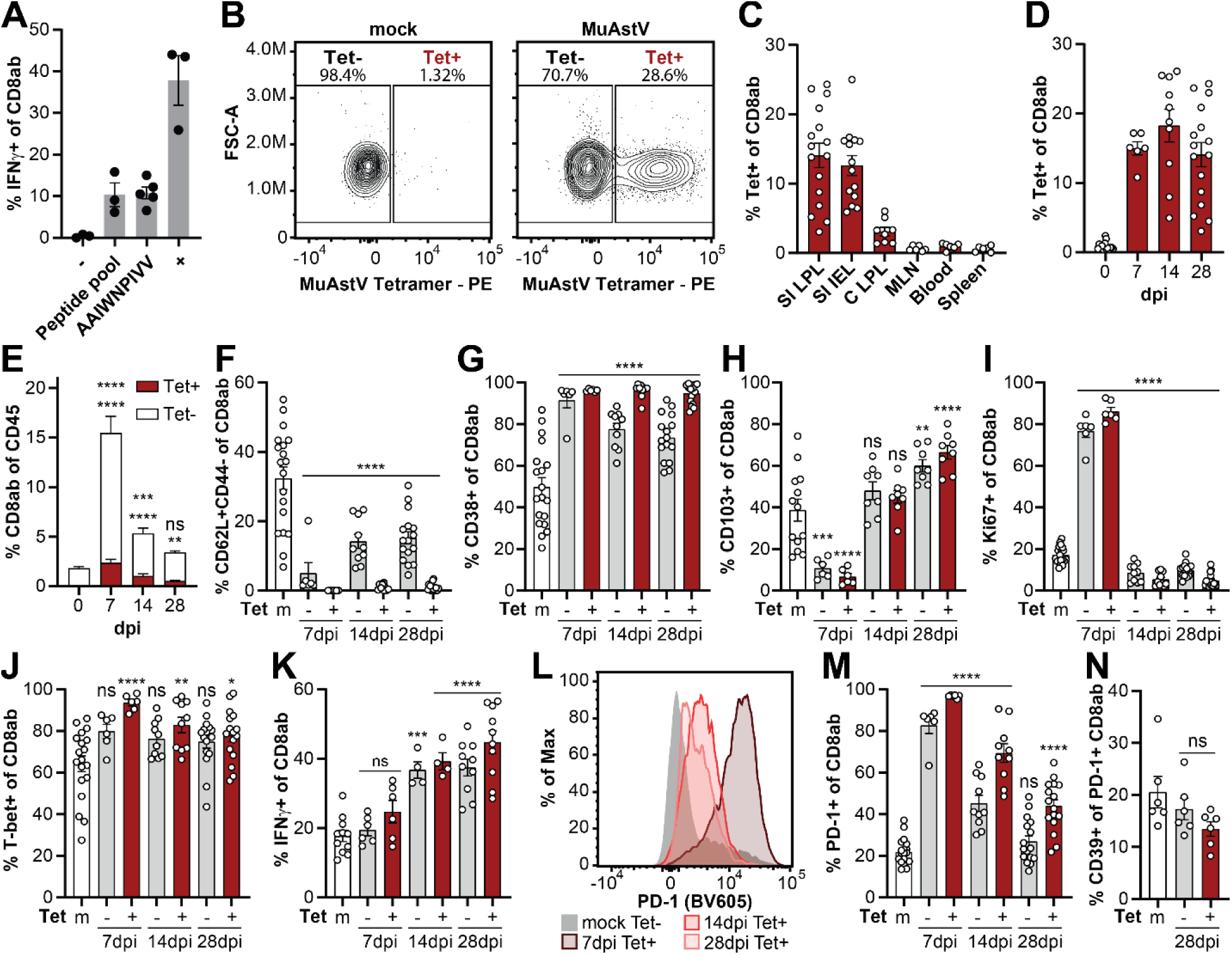
Virus-specific CD8 T cells are not exhausted despite persistent infection: **(A)** Percentage of IFNγ+ CD8 T cells in SI LPL of MuAstV infected mice after stimulation with DMSO (-), a small peptide pool of MuAstV capsid peptides, the immunodominant peptide AAIWNPIVV, or positive control cell stimulation cocktail (+) as measured by flow cytometry. n = 3-4 mice. **(B)** Representative flow plots of MuAstV Tetramer staining of CD8ab T cells in SI LPL for mock and MuAstV mice. Tet-, Tetramer negative; Tet+, Tetramer positive. **(C)** Percentage of Tet+ CD8ab T cells in different organs around 28dpi detected by flow cytometry. **(D)** Percentage of Tet+ CD8ab T cells in SI LPL over time. **(E)** Percentage of Tet- and Tet+ CD8ab T cells out of CD45+ immune cells in SI LPL over time. **(F-K)** Percentage of Tet- and Tet+ CD8ab T cells in SI LPL over time that are (F) CD62L+CD44-, (G) CD38+, (H) CD103+, (I) Ki67+, (J) T-bet+, and (K) IFNγ+, that last after stimulation. **(L)** Representative histograms of PD-1 levels on Tet- and Tet+ populations at different times post infection. **(M-N)** Percentage of Tet- and Tet+ CD8ab T cells that are PD-1+ (M) or CD39+ (N). n = 6-16 mice in (C-N). SPF mice were used for all experiments. Each dot represents data from one animal in panels (A), (C-K) and (M-N). All data are represented as mean ± SEM from at least two independent experiments. ns = not significant, *p≤0.05, **p≤0.01, ***p≤0.001, ****p≤0.0001 by 2way ANOVA and Holm-Šídák’s comparison. See also Figure S3.

A tetramer created using this immunodominant peptide stained up to 30% of CD8 T cells in the small intestine LPL fraction of infected mice (Tet+) by flow cytometry with minimal background signal in mock mice (**Figure 4B and S3E**). We performed the initial characterization using infected SPF mice without cage change to establish a baseline. At 28dpi, ∼15% of CD8 T cells were Tet+ in the LPL and intraepithelial lymphocyte (IEL) fractions of the small intestine (**Figure 4C**). Tet+ CD8 T cells were also found, to a lesser extent, in colon LPL, and negligible numbers were detected in mesenteric lymph nodes (MLN), blood, and spleen at this time point. When we performed a time course analysis focusing on the small intestine LPL and IEL, Tet+ cells were already detectable at 7dpi and remained at ∼15% out of all CD8 T cells in both compartments (**Figure 4D and S3F**). Tetramer negative (Tet-) and Tet+ CD8 T cells represented a large proportion of total CD45+ cells (leukocytes) at 7dpi and then contracted sharply (**Figure 4E and S3G**). The proportion of Tet- and Tet+ cells displaying markers of naïve CD8 T cells (CD62L+CD44-) were low on 7dpi compared with CD8 T cells in mock infected mice. At later time points, many of the Tet-cells were CD62L+CD44- while the Tet+ CD8 T cells with these markers remained low, as expected for antigen-specific cells (**Figure 4F and S3H**). Conversely, the proportion of activated (CD38+) CD8 T cells increased to almost 100% at 7dpi and was stable in the Tet+ fraction (**Figure 4G and S3I**). These results show that MuAstV infection elicits an antigen-specific CD8 T cell response.

The proportions of Tet+ and Tet- CD8 T cells in the LPL and IEL fractions that were positive for the residency marker CD103 decreased at 7dpi, which was followed by a restoration or increase at 14dpi and 28dpi (**Figure 4H and S3J**). In contrast, the proportion of Ki67+ CD8 T cells spiked at 7dpi and then decreased below mock levels by 14dpi (**Figure 4I and S3K**), matching the contraction of CD8 T cells. Most CD8 T cells were positive for the type 1 transcription factor T-bet, and many produced IFNγ, especially Tet+ CD8 T cells at later time points (**Figure 4J-K, S3L-M**). These results are consistent with an influx of CD8 T cells to the small intestine in response to infection that initially proliferate, and then are retained as tissue resident effector cells.

Subsequently, we investigated whether T cells become exhausted, as seen with LCMV^20^. We examined PD-1, an inhibitory marker upregulated after activation and whose failure to be downregulated is a pre-requisite to exhaustion^43^. As expected, PD-1 levels were upregulated at the peak of CD8 T cell response at 7dpi (**Figure 4L+M and S3N**). Afterwards, PD-1 was downregulated, especially in the IEL fraction and the Tet- population in the LPL. As Tet+ cells in the LPL retained higher levels of PD-1, we stained for CD39 as a marker of terminal exhaustion. CD39 levels at 28dpi were not elevated in Tet- and Tet+ cells compared to mock (**Figure 4N**). This data is consistent with CD8 T cells actively reacting to MuAstV at peak of infection followed by T cell ignorance, potentially similar to what has been observed with MNV^24,25^. Furthermore, this suggests that functional virus-specific CD8 T cells are present and ready to react to any disturbance before cage change.

### Cage change evokes CD8 T cell and epithelial defense responses

Flow cytometric analyses of CD8 T cells in LPL and IEL of MuAstV-infected mice 1 day post cage change did not reveal any significant differences in either Tet- and Tet+ populations when compared to the no cage change control, although there was a trend towards decreased naïve and increased replicating CD8 T cells after cage change (**Figure S4A-B**). To examine cellular responses not captured by our flow cytometry analysis, we performed single cell RNA sequencing (scRNA-Seq) of small intestinal tissue from mock- and MuAstV-infected mice with and without cage change focusing on 1 day after the perturbation (**Figure S4C-D**).

Our initial analysis focused on immune cells (CD45+), yielding 15 distinct populations annotated using signatures of known lineage markers (**Figure 5A and S4E-F**). Cytotoxic CD8 T cells were increased in MuAstV mice with cage change (**Figure 5B and S4G**), supporting our flow cytometry data. Further sub-clustering of T cells revealed that CD8 central memory (TCM) and CD8 effector memory (TEM) cells were highest in the MuAstV infection with cage change group (**Figure S4H-J**). Pathway analysis for all T cells revealed that cage change, in both mock and infected mice, resulted in upregulation of genes involved in RNA processing, organelle organization, and intra-cellular trafficking (**Figure S5A**), pathways that have been linked to either T cell activation or formation of immunological synapses^44,45^. Similar pathways were enriched when we analyzed CD8 TEM specifically (**Figure 5C**). CD8 TEM in infected mice also upregulated genes involved in *viral processes* after cage change, containing transcripts with known antiviral roles such as *Ddx5* or *Atg16l1*^46-49^.

**Figure 5.**
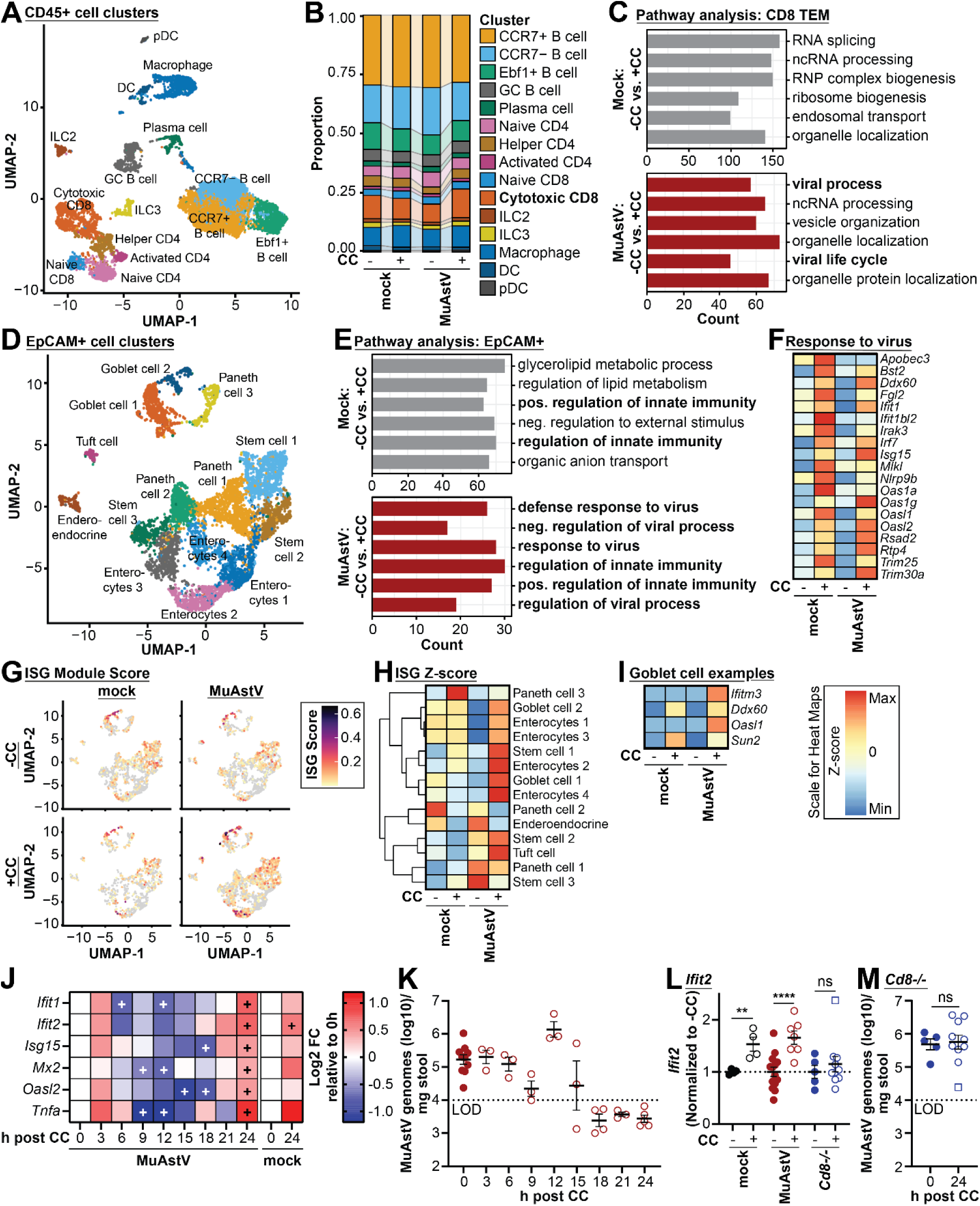
Cage change evokes CD8 T cell and epithelial defense responses: **(A)** UMAP visualization of major immune cell clusters in scRNA-Seq data combining all conditions. Cell from 2 mice were pooled per condition. **(B)** Proportion of immune cell clusters shown in A in each condition. **(C)** Pathway enrichment analysis of genes upregulated in both cage change conditions showing the six most upregulated pathways in mock (top) and MuAstV (bottom) in the CD8 TEM cluster with gene counts for each. **(D)** UMAP visualization of major epithelial cell clusters in scRNA-Seq data. **(E)** Pathway enrichment analysis of genes upregulated in both cage change conditions showing the six most upregulated pathways in mock (top) and MuAstV (bottom) in the EpCAM+ cells with gene counts for each. **(F)** Heat map for z-scores of genes in the ‘response to virus’ pathway in EpCAM+ cells. **(G)** Module Score of the ISG signature in epithelial cell clusters in the individual samples. **(H)** Z-score for ISGs in different EpCAM+ clusters. **(I)** Expression levels of individual ISGs in goblet cells. **(J)** Heat map showing changes in gene expression over time in ileal epithelial layer. Values are normalized to 0h post CC. FC fold change. n = 3-10 mice. **(K)** MuAstV levels from mice in (J). n = 3-10 mice. **(L)** Normalized *Ifit2* RNA levels in mock infected WT, MuAstV infected WT and MuAstV infected *Cd8^-/-^* mice +/-CC. n = 4-14 mice. **(M)** MuAstV levels from *Cd8^-/-^* mice in L. n = 5-10 mice. Square (□) in (L+M) indicates the same mouse that behaved as an outlier. SPF mice were used for all experiments. Each dot represents data from one animal in panels. (K-M) and the data are represented as mean ± SEM from at least two independent experiments. ns = not significant, *p≤0.05, **p≤0.01, ***p≤0.001, ****p≤0.0001 by two-sided Student’s T-test compared to no CC for panels (J) and (L-M). For panel J ‘+’ denotes a fold change with any p-value less than 0.05. See also Figure S4-6.

The EpCAM+ fraction contained the expected epithelial cell populations (**Figure 5D and S5B-C**). Pathway analysis revealed that cage change increased expression of genes related to *innate immune responses* in the epithelium of both mock and infected mice (**Figure 5E and S5D**). Transcripts and pathways associated with antiviral immunity were upregulated upon cage change, especially in infected mice. This category contains interferon-stimulated genes (ISGs), known antiviral factors, and other genes involved in host defense such as those that encode TRIM proteins (**Figure 5F)**. An ISG score, representing the average level of 16 ISGs, showed that intestinal epithelial cells express ISGs at baseline, which was increased the most in infected mice after cage change (**Figure 5G**). Goblet cells, which harbor MuAstV (**Figure 1D**), were among the cell subsets with the largest upregulation of ISGs after cage change (**Figure 5H**). When examining specific antiviral transcripts in goblet cells, we noticed that some were induced upon cage change in both mock and infected mice (e.g., *Ddx60* and *Sun2*), while others were only induced in the presence of MuAstV (e.g., *Ifitm3* and *Oasl1*) (**Figure 5I**). In addition, we observed that goblet cells upregulated *Tnfa*, encoding the inflammatory mediator TNFα. These results show that host defense transcripts are induced in the intestinal epithelium following cage change, which is further enhanced in infected mice.

To examine the timing of these responses, we measured the expression of these defense related genes in the intestinal epithelium at 3h intervals following cage change in MuAstV-infected mice (**Figure 5J**). We observed a modest increase at 3h, followed by a sharp decrease that was lowest at 12h and another increase by 24h post cage change. This second increase in ISG and *Tnfa* expression at 24h was also observed in mock mice but not as strongly (**Figure 5J and S6A-E**). MuAstV levels were increased at 12h post cage change, consistent with the decrease in defense gene expression at that time, followed by a rapid decline (**Figure 5K**).

We hypothesized that CD8 T cells are upstream of this epithelial immune response. We therefore assessed defense gene expression after cage change in *Cd8^-/-^* mice. Most of the genes, with exception of *Isg15*, were not significantly increased in *Cd8^-/-^* mice after cage change (**Figure 5L and S6A-E**). There was one out of ten mice, however, that had very high ISG induction after 24h in a new cage. This heterogenous response is reminiscent of our earlier finding showing that a subset of *Cd8^-/-^* mice retain the ability to clear the virus upon cage change (**Figure 3J-K**). Indeed, *Cd8^-/-^* mice that did not upregulate host defense genes in the epithelium upon cage change displayed high levels of MuAstV infection while the one mouse with successful induction of these transcripts showed a reduction in vRNA (**Figure 5M**).

Together, these data indicate that cage change results in immune activation in both the immune and epithelial compartment, which is enhanced in the presence of MuAstV. When the CD8 T cells successfully inhibit the virus, their activity lies upstream of an antiviral response by the epithelium.

### Glucocorticoid fluctuation is necessary for host defense and virus clearance

The initial decrease in defense gene expression at 12h post cage change was unexpected. Among pathways enriched after cage change in both mock and infected mice was *response to peptide hormones* (**Figure S5D**). Hormones released upon stress, such as glucocorticoids, have potent anti-inflammatory functions^50^. We found that several genes that are affected by glucocorticoid signaling were responsive to cage change (**Figure 6A**). We examined the expression of one gene upregulated by glucocorticoids (*Sgk1*) at additional timepoints (**Figure 6B**). *Sgk1* exhibited a short-term amplification at 3h and then a peak at 12h, followed by a sharp decline at 18h and returning to baseline by 24h. Serum levels of corticosterone, the main murine glucocorticoid, increased up to 12h post cage change and then dropped sharply, matching the expression pattern of *Sgk1* (**Figure 6C**). This initial increase and then drop in corticosterone corresponded remarkably well with an inverse pattern of defense gene expression that initially decreased and then rebounded to above baseline levels (**Figure 6D**). Overlaying the time course of MuAstV levels on top of these measurements further revealed that the corticosterone-associated dip in defense gene expression was followed by a spike in virus levels and then a subsequent sharp decline as corticosterone and defense gene expression decreased and increased, respectively (**Figure 6D**).

**Figure 6.**
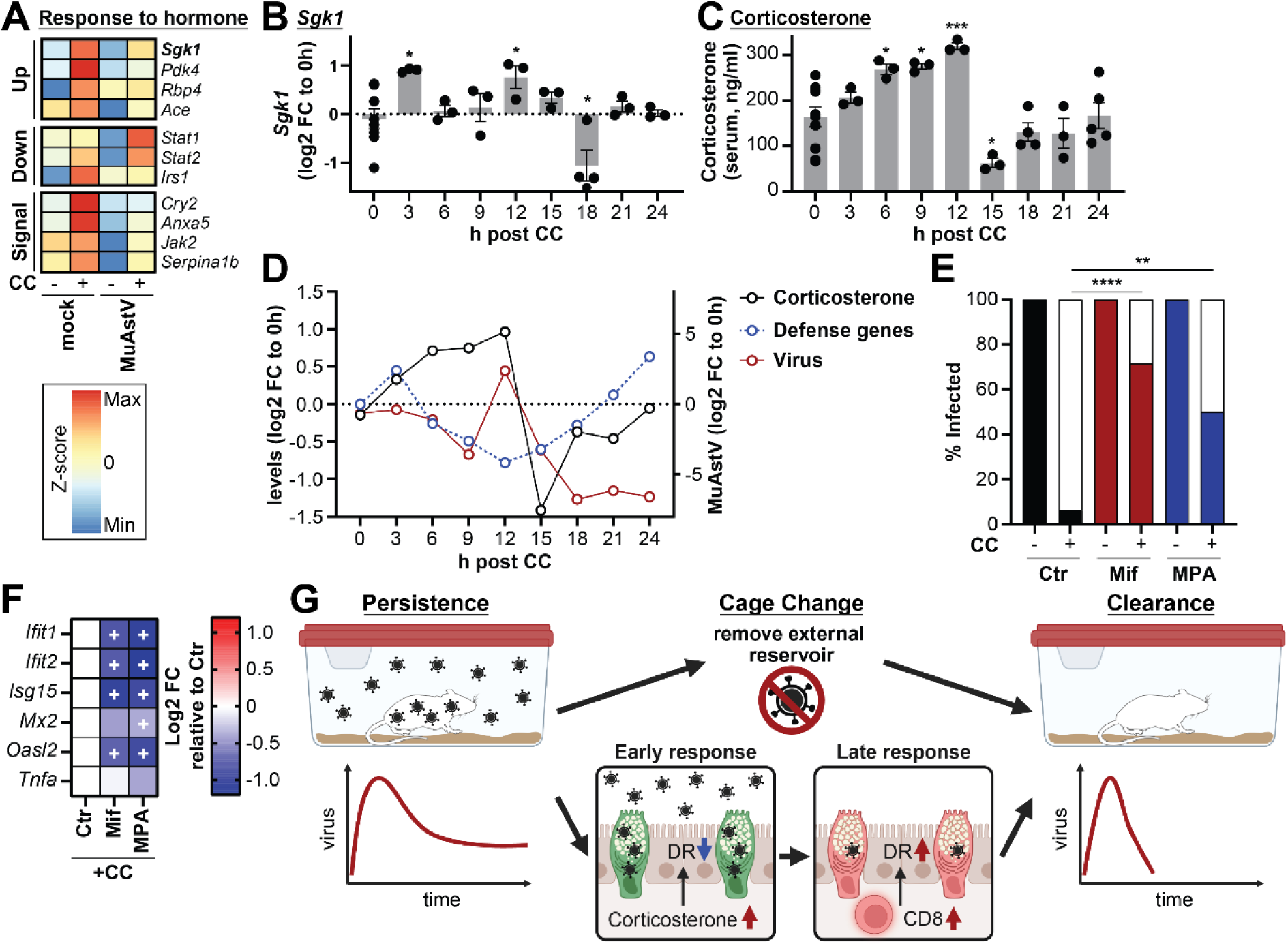
Glucocorticoid fluctuation is necessary for ISG induction and virus clearance: **(A)** Heat map for z-scores of genes in the *response to peptide hormone* pathway that are related to glucocorticoid signaling in EpCAM+ cells in scRNA-Seq dataset. Genes are grouped based on whether prior studies have shown them to be upregulated (Up), downregulated (Down), or interact with the glucocorticoid receptor during hormone signaling (Signal). **(B)** Time course of *Sgk1* expression after cage change in samples from Fig. 5J. n = 3-8 mice. **(C)** Corticosterone levels in serum from mice from Fig. 5J as determined by ELISA. n = 3-10 mice. **(D)** Comparison of behavior of averaged corticosterone levels, defense gene expression, and MuAstV levels after cage change. **(E)** Percentage of mice with detectable MuAstV in their stool treated with either vehicle control (Ctr), glucocorticoid antagonist Mifepristone (Mif) or synthetic glucocorticoid Methylprednisolone acetate (MPA) +/-CC. n = 6-11 mice. **(F)** Defense gene expression in Mifepristone and Methylprednisolone acetate treated mice 24h post cage change compared to control. n = 4-6 mice. **(G)** Model of cage change-induced MuAstV clearance. DR = defense response. SPF mice were used for all experiments. Each dot represents data from one animal in panels (B-C) and (E-F) and the data are represented as mean ± SEM from at least two independent experiments. ns = not significant, *p≤0.05, **p≤0.01, ***p≤0.001, ****p≤0.0001 by 2way ANOVA using Holm-Šídák’s comparison for panels (C) and (E), and two-sided Student’s T-test compared to Ctr for panel (F). For panel F ‘+’ denotes a fold change with any p-value less than 0.05. See also Figure S6.

We hypothesized that the short-lived anti-inflammatory glucocorticoid signaling following cage change was leading to a rebound reaction by the immune system that mediated MuAstV clearance. If the rebound reaction is necessary, we predicted that breaking this pattern by either blocking glucocorticoid signaling or constitutively activating the pathway should both interfere with cage change-mediated viral clearance. Indeed, administration of the antagonist Mifepristone or the synthetic glucocorticoid Methylprednisolone acetate significantly inhibited MuAstV clearance after cage change (**Figure 6E**). In addition, defense gene expression was dramatically reduced by both drugs in the context of cage change (**Figure 6F**), placing glucocorticoids upstream of defense gene induction.

Altogether, these data demonstrate stress hormones released upon cage change play an important role in initially suppressing intestinal immunity, followed by increased virus release, defense gene induction, and ultimately viral clearance.

## Discussion

We discovered that an intervention as simple as changing cages determines the duration of a model viral infection in mice. Our data support the following sequences of events leading to MuAstV clearance. Cage change causes a stress response characterized by immediate and temporary production of corticosterone. This corticosterone initially triggers immune suppression and a transient increase in viral replication. Once recovered from the immune suppression, there is a rebound in intestinal CD8 T cells and a related antiviral defense response in the epithelium (**Figure 6G**). We posit that this rebound leads to a heightened state of immunity above pre-cage change levels that mediates elimination of viruses. This antiviral response is likely not sufficient to prevent reinfection when viruses remain in the same environment, thus explaining why removing the external reservoir and breaking the chain of reinfection is necessary for resolving the infection.

Persistent infections might be more common than appreciated for MuAstV in natural settings. The high degree of prevalence of MuAstV in wild and pet store mice^51-53^ may reflect prolonged infections and windows of transmission. Although free-living mammals do not experience cage changes, predators or natural disasters can force infected animals to migrate under duress. In those cases, the removal of the external reservoir and the stress-mediated strengthening of the intestinal immune barrier could induce clearance of ongoing infections while protecting from potential new pathogens present in the next locale.

Our observations also have implications for mouse husbandry. Cage change during the daytime is standard practice but causes considerable stress by disrupting the normal resting period of mice, which are nocturnal^37^. Despite representing such a drastic procedure, cage changes are rarely considered as an experimental variable in mouse studies. Through personal communications, we have learned of anecdotal examples in which researchers observed unexplained changes in their mouse models of infectious disease and cancer during the laboratory shutdown period of the COVID-19 pandemic when cage changes occurred less frequently due to personnel and supply restrictions. Careful documentation of husbandry practices may aid with efforts to improve experimental reproducibility or reveal pathways that are sensitive to external perturbations.

Stress and glucocorticoids have established immunomodulatory effects and clinical applications, but the target cell types and signaling pathways are frequently not known because they are tissue- and context-specific^54^. Glucocorticoids can impact CD8 T cells by favoring memory over effector programming, which is important for limiting lethal immunopathology during infections^55-57^. Although typically considered anti-inflammatory, sustained glucocorticoid production due to psychological stress exacerbates TNFα-induced intestinal inflammation^58^. In our model, this counterintuitive enhancement of *Tnfa* expression and other host defense genes was explained by timing and best described as a rebound effect when glucocorticoids levels sink below baseline levels. Given the variety of external and internal factors that can trigger a stress response, an important future direction will be to define the exact relationship between corticosterone, CD8 T cells, and the epithelium in a temporal manner, and determine whether this axis applies to other organs.

Glucocorticoid administration during the resolution of a viral infection improves health outcomes by preventing immune-mediated damage, which is especially important when antivirals are unavailable^59^. Our results suggest that persistence of the virus may be impacted by the timing of such therapy. Additionally, the degree of psychological stress and opportunities for re-exposure to a virus can vary considerably depending on whether an infected individual is symptomatic, admitted to a hospital, or sharing living quarters with others who are infectious such as family members. These variables potentially explain why a growing list of viruses or their products can be detected in a subset of individuals but not others after recovery from acute illness^17^. A better understanding of how external factors impact immune responses may reveal non-pharmaceutical intervention strategies or inform the use of existing drugs to control the duration of a viral infection.

## Figures

**Figure S1:**
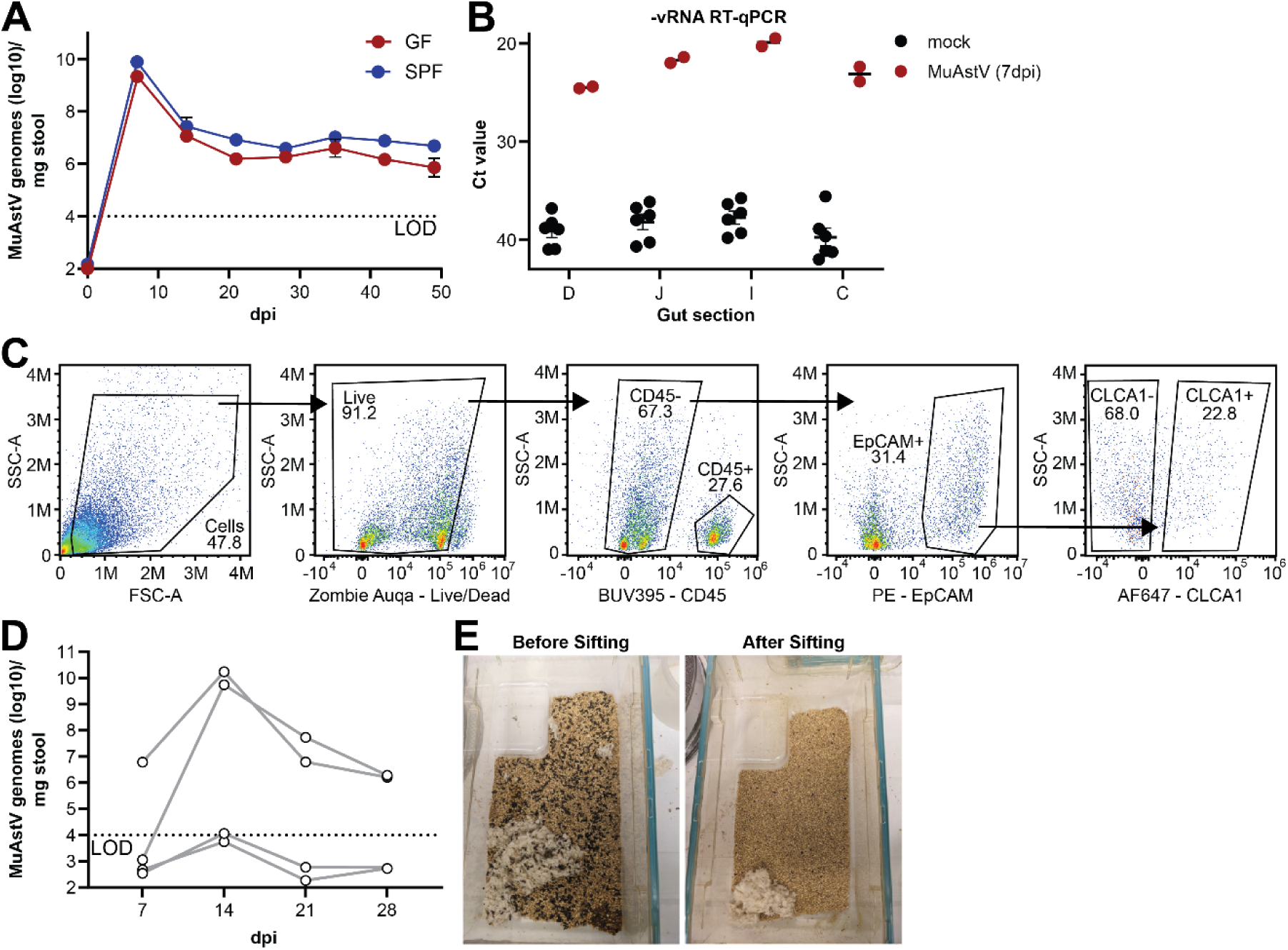
**(A)** MuAstV genome levels in stool of GF versus SPF B6 mice over time without CC detected by RT-qPCR. n = 3 mice. **(B)** Ct values for MuAstV negative strand RT-qPCR mock mice and those at peak of infection at 7dpi in different gut sections. D - Duodenum, J - Jejunum, I - Ileum, C - Colon. Each dot represents data from one animal. n = 2 mice. **(C)** Representative flow plots illustrating the gating strategy used for sorting of the following population in Figure 1E: Immune - CD45+, Epithelial (-GC) - EpCAM+CLCA1-, Goblet Cells - EpCAM+CLCA1+. **(D)** MuAstV levels for individual mice infected with 1E6 viral genomes. n = 4 mice. **(E)** Representative images of cages before and after removal of the environmental viral reservoir through sifting. GF mice used in panel (A), SPF used in all other panels. Data in (A) are represented as mean ± SEM.

**Figure S2:**
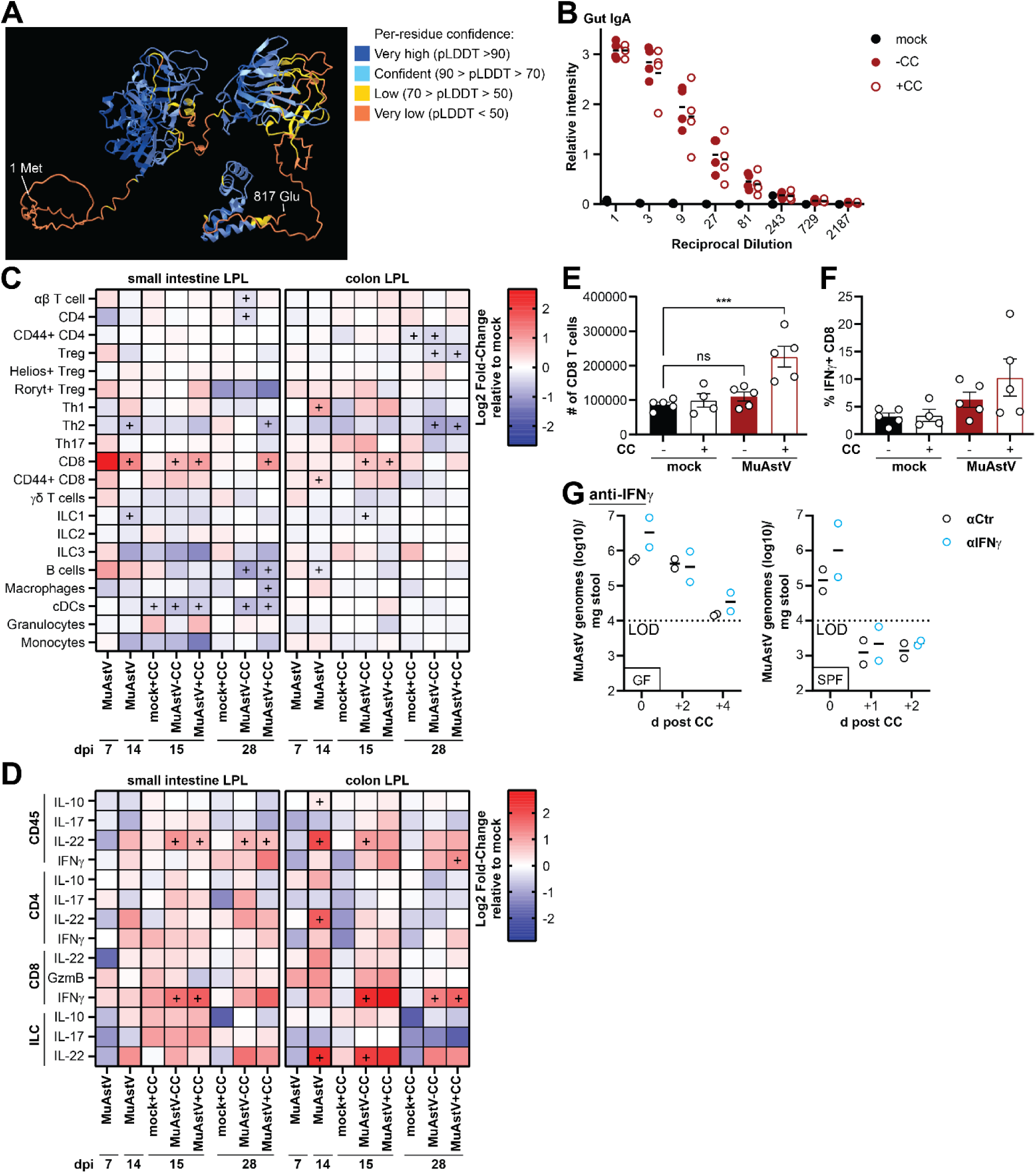
**(A)** 3D structure of MuAstV Orf2 as predicted with AlphaFold and corresponding confidence. Amino acids 425-686 are the spike portion of the MuAstV capsid and represent the immunodominant domain for humoral responses. **(B)** Raw values for anti-spike IgA data shown in Figure 3B. n = 4 mice. **(C-D)** Heatmaps of average fold changes compared to mock-CC for immune populations (C) or cytokine production (D) in the small intestine (SI) and colon (C) as frequencies of LPL in GF mice over time as measured by flow cytometry. n = 2-6 mice. **(E)** Total number of CD8 T cells in the SI LPL fraction at 28dpi. n = 4-6 mice. **(F)** Percentage of CD8 T cells producing IFNγ after stimulation at 28dpi. n = 4-6 mice. **(G)** MuAstV levels after CC in GF or SPF mice that were either pre-treated with a control or anti-IFNγ antibody. n = 2 mice. GF mice used in panel (B-F), data for GF and SPF mice pooled for panel (G). Each dot represents data from one animal in panel (B) and (E-G). Data in (E-F) are represented as mean ± SEM. ns = not significant, **p≤0.01, ***p≤0.001, ****p≤0.0001 by 2way ANOVA and Holm-Šídák’s comparison for (E), and two-sided Student’s T-test for (B-C). For panel (C-D) ‘+’ only denotes a fold change with any p-value less than 0.05.

**Figure S3:**
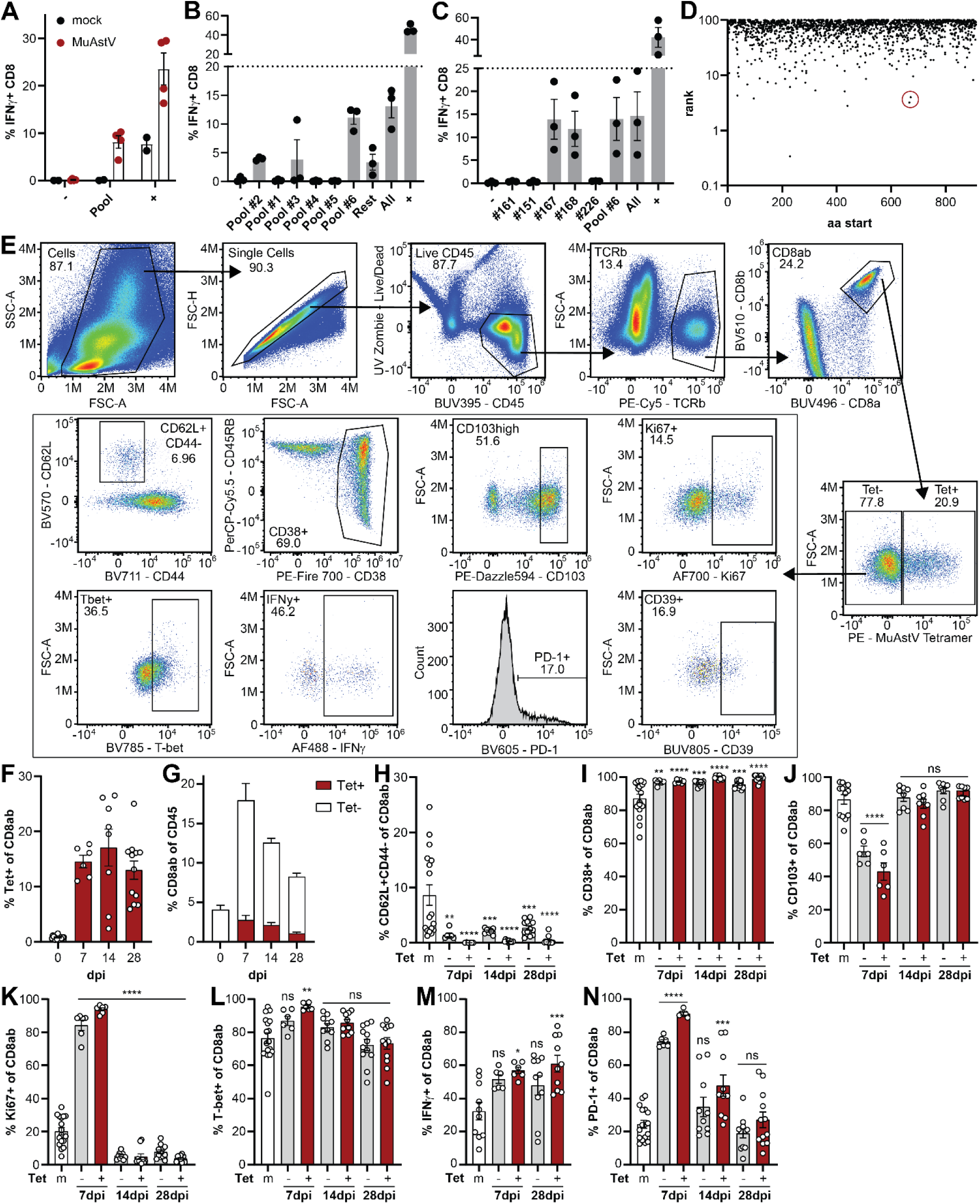
**(A)** Percentage of IFNγ+ CD8 T cells in SI LPL of mock or MuAstV mice after 5h of stimulation with either DMSO (-), a small peptide pool of MuAstV capsid peptides containing the 50 peptides with the highest binding capacity to mouse MHC I/II (pool), or cell stimulation cocktail (+) as measured by flow cytometry. n = 2-4 mice. **(B)** Percentage of IFNγ+ CD8 T cells in SI LPL of MuAstV mice stimulated with smaller peptide pools or controls. n = 3 mice. **(C)** Percentage of IFNγ+ CD8 T cells in SI LPL of MuAstV mice stimulated with individual peptides or controls. n = 3 mice. **(D)** Rank of MHC I binding prediction of MuAstV Orf2 capsid protein. Identified immunodominant peptides are highlighted with the red circle. **(E)** Representative flow plots illustrating the gating strategy using the MuAstV-specific Tetramer to characterize CD8 T cell responses after viral infection. **(F)** Percentage of Tet+ CD8ab T cells in SI IEL over time. **(G)** Percentage of Tet- and Tet+ CD8ab T cells out of CD45+ immune cells in SI IEL over time. **(H-N)** Percentage of Tet- and Tet+ CD8ab T cells in SI IEL over time that are (H) CD62L+CD44-, (I) CD38+, (J) CD103+, (K) Ki67+, (L) T-bet+, (M) IFNγ+ after stimulation and (N) PD-1+. n = 6-16 mice in (F-N). Each dot represents data from one animal in panels (A-C) and (F-N). All data are represented as mean ± SEM from at least two independent experiments. ns = not significant, *p≤0.05, **p≤0.01, ***p≤0.001, ****p≤0.0001 by 2way ANOVA and Holm-Šídák’s comparison.

**Figure S4:**
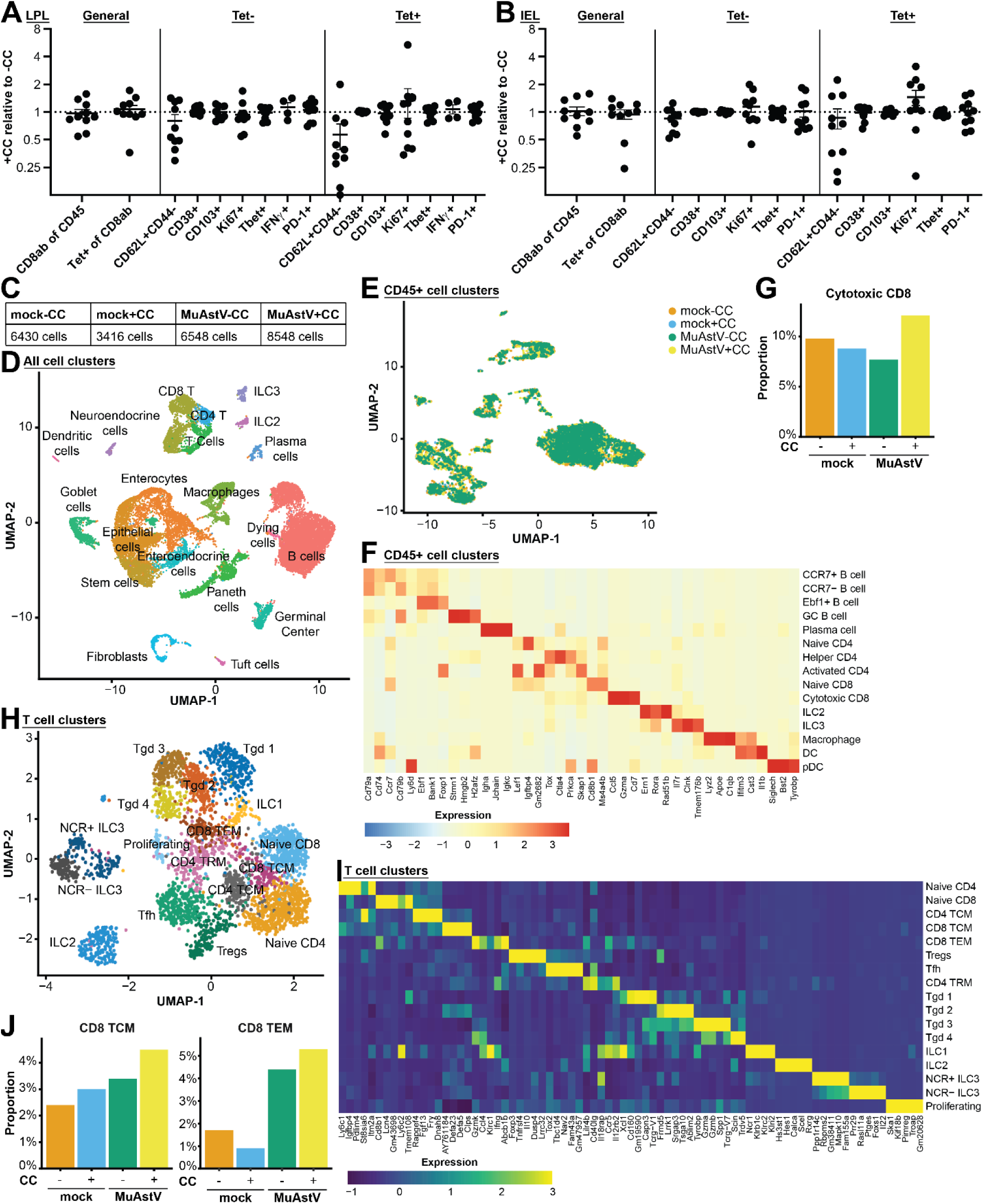
**(A-B)** Comparison of Tet- and Tet+ populations in the SI LPL (A) and IEL (B) in MuAstV infected mice 1d post CC as compared to the average of the -CC controls. Each dot represents one animal. Data are represented as mean ± SEM from at least two different experiments. Statistical analysis via two-sided Student’s T-test revealed no significant differences. n = 4-10 mice. **(C)** Cell numbers for singe-cell RNA sequencing (scRNA-Seq) for the different conditions that passed quality control. **(D)** UMAP visualization of all cells in the scRNA-Seq data set. **(E)** UMAP visualization of CD45+ cells in the data set color coded by sample. **(F)** Heatmap of lineage marker expression used for defining the different immune cell clusters. **(G)** Proportion of cytotoxic CD8 under the different conditions in the immune cell data set. **(H)** UMAP visualization of the T cell subclusters. **(I)** Heatmap of lineage marker expression used to define the different T cell subclusters. **(J)** Proportion of CD8 TCM and TEM under the different conditions in the T cell subset.

**Figure S5:**
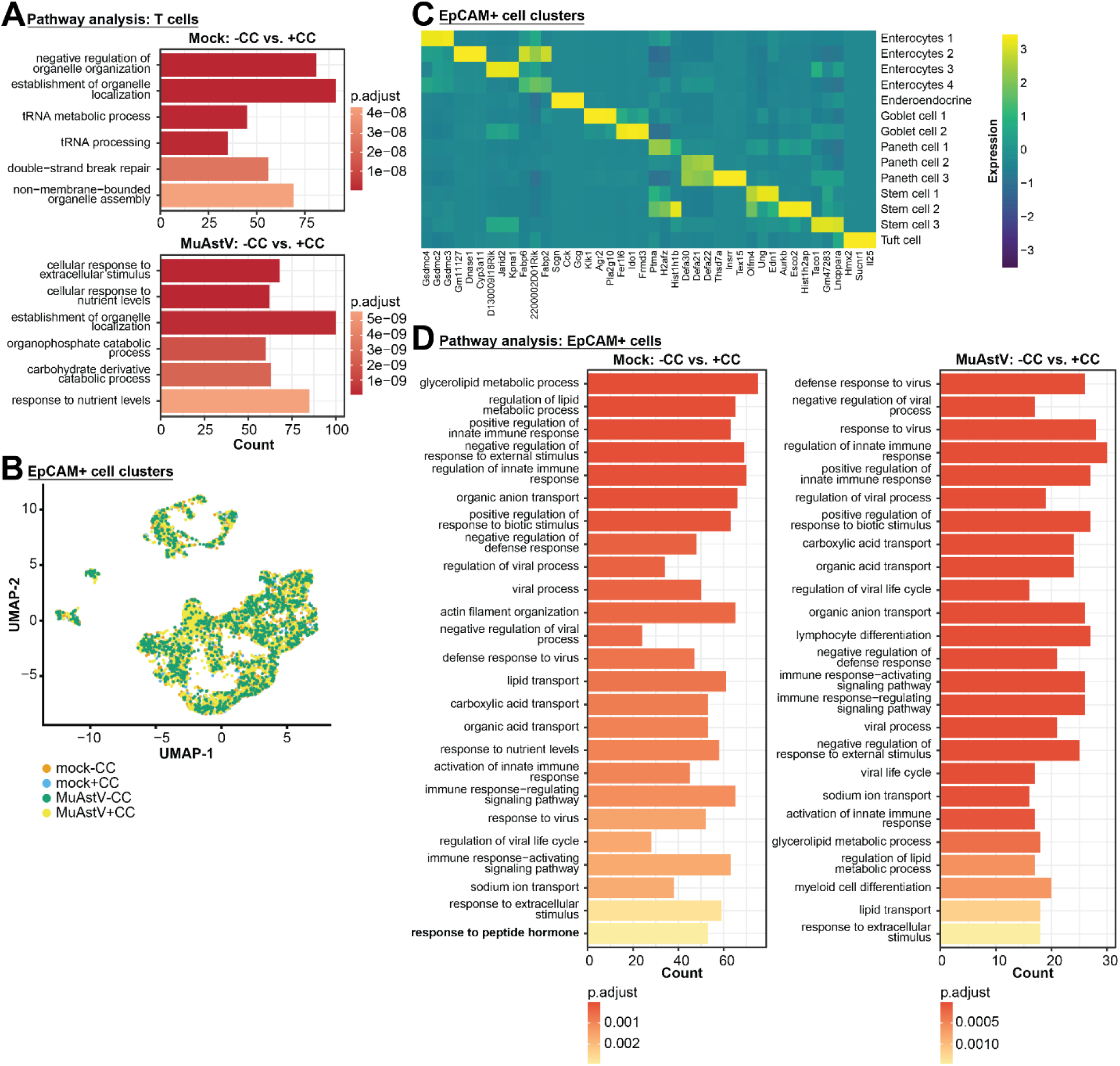
**(A)** Pathway enrichment analysis of genes upregulated in both cage change conditions showing the six most upregulated pathways in mock (top) and MuAstV (bottom) in the T cell subset with gene counts for each. **(B)** UMAP visualization of EpCAM+ cells in the data set color coded by sample. **(C)** Heatmap of lineage marker expression used for defining the different epithelial cell clusters. **(D)** Pathway enrichment analysis of genes upregulated in both cage change conditions showing the six most upregulated pathways in mock (left) and MuAstV (right) in the EpCAM+ subset with gene counts for each.

**Figure S6:**
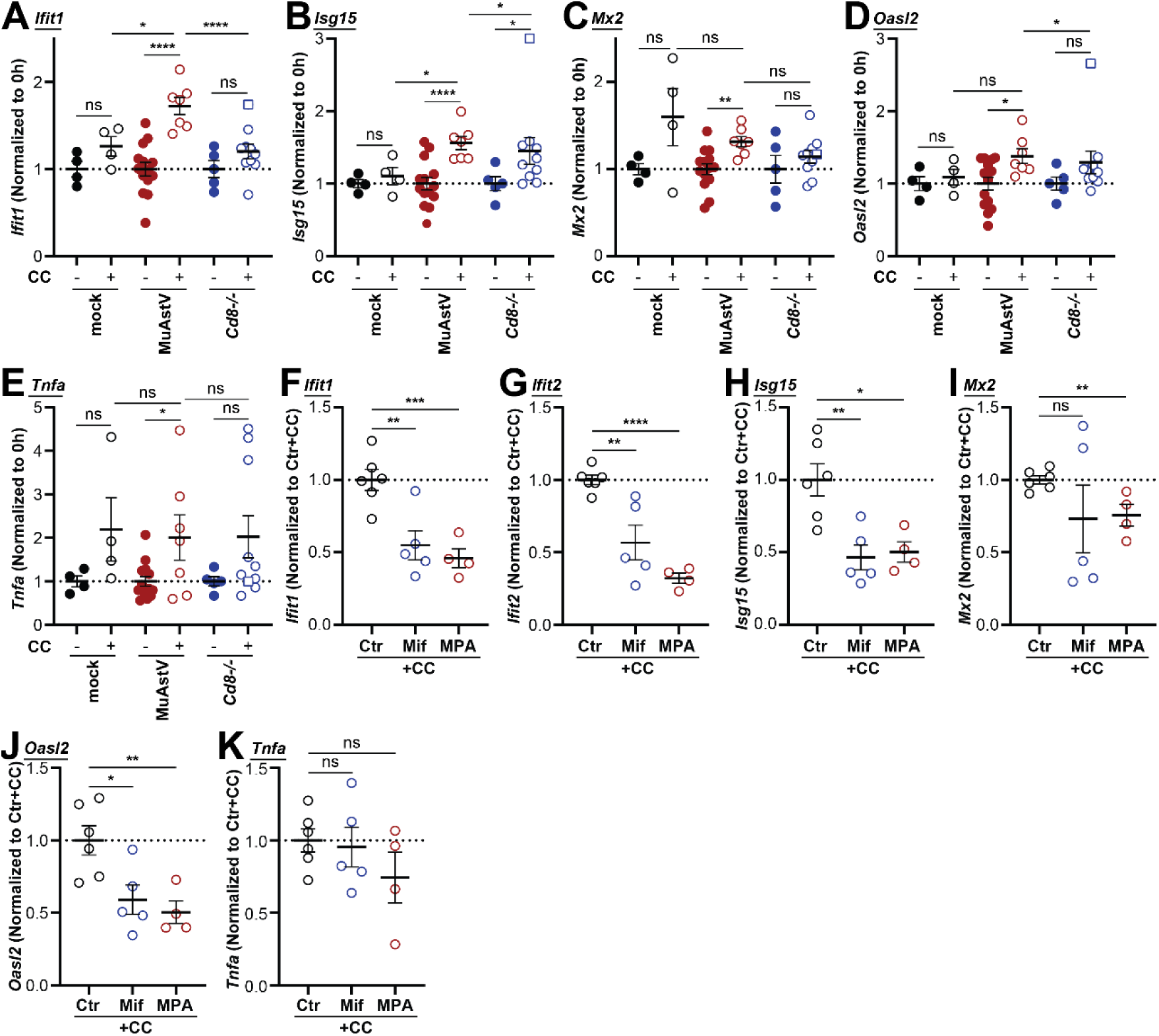
**(A-E)** Normalized RNA levels in mock infected WT, MuAstV infected WT and MuAstV infected CD8 KO mice +/-CC for the following genes: (A) *Ifit1*, (B) *Isg15*, (C) *Mx2*, (D) *Oasl2*, and (E) *Tnfa*. Square (□) indicates the same mouse that behaved as an outlier and was excluded from statistics. n = 4-14 mice. **(F-K)** Normalized RNA levels in control, Mifepristone and Methylprednisolone acetate treated mice 24h post cage change for the following genes: (F) *Ifit1*, (G) *Ifit2*, (H) *Isg15*, (I) *Mx2*, (J) *Oasl2*, and (K) *Tnfa*. n = 4-6 mice. In all the panels each dot represents one animal. Data are represented as mean ± SEM from at least two different experiments. ns = not significant, *p≤0.05, **p≤0.01, ***p≤0.001, ****p≤0.0001 via two-sided Student’s T-test in all panels.

## Methods

### Mice

C57BL/6J (WT, #000664), *Cd8^-/-^* (#002665) and *Ifngr^-/-^* (#003288) were originally purchased from The Jackson Laboratory and bred onsite or directly used for experiments. *Rag1^-/-^* mice were obtained from the Gnotobiotic Mouse Core at the University of Pennsylvania to avoid MuAstV contamination risk. Mice were confirmed to be negative for MuAstV by RT-qPCR before experiments. For mice that were positive for MuAstV infection, the line underwent backcrossing through C57BL/6J or early cross-fostering to eliminate virus contamination.

Mice between 6-12 weeks of age were used for all experiments. Age- and sex-matched littermate controls were used whenever possible. Mice were housed in ventilated cages at room temperature with a 12h light-dark cycle and given food and water ad libitum. Mice of both sexes were used for all experiments besides the immune profiling in Figure 3 and the scRNA-Seq in Figure 5, for which only female mice were used. GF mice were bred in flexible-film isolators either in the New York University Grossman School of Medicine Gnotobiotics Animal Facility or the University of Pennsylvania Gnotobiotic Mouse Core. For experiments, GF mice were transferred to ISOcages (Tecniplast) with sterile food and water, and absence of bacteria was monitored throughout the experiment by qPCR for 16S. All experiments were performed in accordance with IACUC protocols approved by the New York University or University of Pennsylvania.

Cage change involved transfer of mice into a clean cage with fresh food and water. Cage trade involved transfer of mice into a cage that previously housed other mice of the same sex. Cage sifting was performed by transferring mice to a clean cage temporarily (up to 15 minutes) and removing as much feces as possible by sifting the cage contents with a Garden Potting Mix Sieve (Practicool) by combining the 2, 3 and 4mm mesh filters followed by manual removal of remaining stool pellets. For viral passage, infected mice were housed with uninfected mice at 14dpi. A pilot study was conducted for prolonged housing of mice without cage change (manuscript in preparation). In short, ammonia levels, humidity and temperature were measured daily for 12 weeks. It was determined that housing of 2 mice in the same cage for up to 6 weeks was possible without increase of ammonia above 50 ppm. Food and water were replenished as needed.

### Virus purification and infection

We used murine astrovirus-1 strain NYU1 for all experiments^42^. Viral stock was generated as previously described^42^. In short, GF C57BL/6J were inoculated and fecal pellets containing virus collected at peak of infection before onset of the humoral immune response. This was followed by homogenization in PBS, removal of particulates by repeated spins at 17000g and filtration of the supernatant through 0.22μm Millex-GP syringe-driven filter unit (Millipore Sigma) at least twice. Viral titers were determined by RNA extraction and RT-qPCR for the viral genome. Mice were inoculated with a standard dose of 1E10 genome copies/mouse of MuAstV diluted in PBS via oral gavage. For repeated gavage of mice with virus, stool pellets from 7dpi of the respective mice were used and equal to 1 pellet/mouse/time point was homogenized in PBS and administered by oral gavage. For the neutralization assay, 10µl of virus was incubated with 40µl of serum for 90min at RT. Virus was diluted 1:100 in PBS for inoculation of mice equaling the standard infectious dose used.

### Viral quantification

Fresh stool pellets were collected from mice before infection and throughout the experiment and stored at -70°C until further processing. Pellets were weighed and homogenized in RLT buffer (RNeasy kit, Qiagen) with 1.0 mm Zirconia/Silica Beads (BioSpec Products) using a FastPrep-24 homogenizer (MP Biomedicals). Debris was removed using two spins of 2,000g and 10,000g, each 5 min at RT. RNA was isolated as described below. The following probes were used for quantifying viral genomes: MuAstV Fwd 5’-TAC ATC GAG CGG GTG GTC GC-3’ and Rev 5’-GTG TCA CTA ACG CGC ACC TTT TCA-3’. Absolute amounts were calculated by comparison to a linearized plasmid.

### Negative strand RT-qPCR

The protocol was adapted from a similar approach used for MNV^60^. RNA was isolated from 1cm of the respective gut sections from mice at 21dpi with and without cage change at 14dpi using the homogenization method mentioned above. 100ng of total RNA were used for reverse transcription using Superscript-IV enzyme (Invitrogen) using the following primers: MuAstV RTnegTag 5’-**GTC CAA CGT GGT GAA GGA TTG TCG** GGA TAA GGC AAG TAG TGC TCC CCT ATT C-3’. The RT primer contained a non-viral tag sequence (bolded) attached to the strand specific sequence. The RT reaction contained 5X buffer, 20mM DTT, 0.5mM dNTPs, 100nM RT-primer and 0.025µl of Superscript-IV enzyme with a total volume of 20µl. The RT reaction was carried out at 55°C for 15min followed by 80°C for 10min. cDNA was diluted 1:10 in H_2_O and qPCR was performed using the following primers: MuAstV neg. strand Fwd 5’-GTC CAA CGT GGT GAA GGA TTG TCG-3’ and Rev 5’-GTA CAA GGA CAT CTT TGG CAT GTG GG-3’.

### RNA isolation and quantification

RNA was isolated using the RNeasy kit including the DNase incubation step (Qiagen). cDNA was generated using High-Capacity cDNA Reverse Transcription Kit (Applied Biosystems) according to the manufacturer’s protocol and stored at −20°C. Transcripts were quantified using either the LightCycler 480 SYBR Green I Master (Roche) on a Roche480II Lightcycler or the PowerTrack™ SYBR Green Master Mix for qPCR (Applied Biosystems) on a QuantStudio 5 Real-Time PCR System (Applied Biosystems).

### MuAstV Spike purification and ELISA

3D structure of the MuAstV NYU-1 capsid protein was modeled using a public version of AlphaFold^38^(https://colab.research.google.com/github/deepmind/alphafold/blob/main/notebooks/AlphaFold.ipynb). Alignment with processing sites reported for human Astrovirus cut sites were used to decide design ELISA epitope (amino acids 425-686)^61^. A gene block covering the spike portion of NYU-1 strain was ordered from IDT and cloned into pGSTag plasmid (Addgene, 21877) through Gibson Assembly (NEB, E2611) using the following primer: Spike-GST Fwd GGT GGT GGT GGT GGA ATT CTA GAC ATG CCT GAA ACC GCA G and Rev G TCA GTC AGT CAC GTG AAT TAA GCT T CTA AGG CCC AGT GGG TGG TAT TTC. Construct was transformed into BL21 (DE3) cells (NEB C2527I) and expression induced with 0.1 mM isopropyl-β-d-thiogalactoside (IPTG; Sigma) for 4h. Cells were lysed (Buffer: 50 mM Tris-HCl pH 8, 10 mM NaCl, 2 mM MgCl2, 1 mM PMSF, Protease inhibitor cocktail, 0.05% NP-40, 0.5 mg/ml Lysozyme and 25 U/ml benzonase), spun for 30min at 4°C 4,000g and protein purified from supernatant using glutathione agarose beads (Pierce, 16100) and elution with reduced L-Glutathione (VWR, 76177-906). Protein purity was confirmed on a protein gel and concentration was determined using a Nanodrop, diluted in 25% glycerol and stored at -20°C.

For the MuAstV ELISA, 96 well plates (USA Scientific, 5667-5074) were coated with 2µg/ml MuAstV spike overnight at 4°C. Wells were washed and blocked in block buffer (PBS, 2%BSA, 0.05% Tween) for 2h at room temperature. After washing, serum samples isolated by cheek bleed or cardiac puncture were diluted in block buffer, added to the plate and incubated for 2h at room temperature. After washing goat anti-mouse IgG or IgA HRP antibody (Invitrogen, 31430) was added for 1h at room temperature. Plate was washed, ELISA developed using TMB (eBioscience, 00-4201-56), stopped using H_2_SO_4_ and read at 450nm using a standard plate reader.

### Corticosterone ELISA

Corticosterone concentrations in serum isolated by cardiac puncture were measured using the Corticosterone Competitive ELISA Kit (Invitrogen, EIACORT) as per manufacturer’s instructions. Mice composing the 0h control group (no cage change) were euthanized at 0h, 11h, 15h and 24h relative to cage change to account for circadian variation in hormone levels.

### Drug administration and other treatments

Mifepristone (MilliporeSigma, M8046) was dissolved in corn oil and administered at 200mg/kg via oral gavage daily from -2 to 0 days relative to cage change. Methylprednisolone acetate (MPA, MilliporeSigma, PHR1718) was dissolved in corn oil and 2mg administered via i.p. once 3 days before cage change. Control mice were administered equal volume of corn oil. Anti-mouse IFNγ antibody (BioXCell, BE0055) or control antibody (BioXCell, BE0088) were administered at 1mg via i.p. at day -3 and 0 relative to cage change.

### Organ processing

For single cell suspensions of small intestinal and colonic LPL fractions, sections were washed with HBSS (Gibco), fat and Peyer’s patches removed, incubated first with 20 mL of HBSS (Gibco) with 2% HEPES (Corning), 1% sodium pyruvate (Corning), 5mM EDTA, and 1 mM dithiothreitol (DTT; Sigma-Aldrich) for 15 min at 37°C and then washed with 10 mL of HBSS with 2% HEPES, 1% sodium pyruvate, 5mM EDTA for 10 min at 37°C. Tissue sections were then washed in HBSS with 5% FCS, minced, and enzymatically digested in HBSS with 5% FCS, collagenase D (0.5 mg/mL, Roche) and DNase I (0.01 mg/mL, Sigma-Aldrich) for 30 min at 37°C with constant agitation. Digested tissue was passed through 70μm mesh and cells were subjected to gradient centrifugation using 40% Percoll (Sigma-Aldrich). For IEL fraction, supernatant after DTT step was washed with RMPI, filtered and subjected to gradient centrifugation using 40% Percoll. MLNs were collected and passed through 70μm cell strainers (BD). Spleens were passed through a 70μm cell strainer, subjected to red blood cell lysis using ACK buffer (Thermo Fisher) and resuspended in PBS. Blood was collected by cardiac puncture into EDTA coated tubes and subjected to red blood cell lysis using ACK buffer (Thermo Fisher) twice and resuspended in PBS.

For measuring cytokine production, cells were plated in RPMI with 10% FCS and 2% HEPES and incubated with Cell Stimulation Cocktail plus protein transport inhibitors (eBioscience) for 4h at 37°C. For goblet cell sorting, about 10cm of the ileum were collected, washed, digested in HBSS with 5% FCS with collagenase D (1 mg/mL) and DNase I (0.15 mg/mL, Sigma-Aldrich) for 30 min at 37°C with constant agitation and filtered through a 70μm cell strainer. For gene expression measurements in the intestinal epithelium, 7.5cm of the ileum were removed and incubated in 10ml of HBSS with 2% HEPES, 1% sodium pyruvate, 5mM EDTA, and 1 mM DTT for 15 min at 37°C. Cells were stored at -70°C until RNA isolation.

### Flow cytometry and cell sorting

Cells were pre-incubated with TruStain FcX™ PLUS (anti-mouse CD16/32) Antibody (BioLegend) and Zombie UV Fixable Viability dye (BioLegend). For Figure 3 three separate staining panels were adapted from Dallari et. al^42^. The first panel included antibodies against CD90.2, CD64, CD11c, CD45, MHCII, B220, CD11b, LY6C and CD103. The second panel included antibodies against CD19, GATA3, Helios, TCRγδ, T-bet, CD3, CD44, NK1.1, CD127, CD62L, CD8β, RORγt, CD45, CD4, CD8α and FoxP3. The third panel included antibodies against CD19, IFNγ, Granzyme B, TCRγδ, IL-22, CD3, IL-17, CD127, TCRβ, IL-10, CD45, CD4 and CD8α. Samples were fixed with either Fixation Buffer (BioLegend) or FoxP3/Transcription Factor Staining Buffer Set (eBioscience). For transcription factors staining cells were permeabilized and stained in the Foxp3/Transcription Factor Staining Buffer Set at 4°C. For cytokine staining cells were permeabilized and stained with the Intracellular Staining Permeabilization Wash Buffer (BioLegend) at 4°C. Samples were recorded on a BD LSR II (BD Biosciences) and analyzed using FlowJo software (FlowJo LLC).

For Figure 5 two separate staining panels were used. These panels included antibodies against CD8α, CD45, CD44, PD-1, CD62L, CD8β, MuAstV Tetramer, TCRβ, CD45RB, CD103, CD38, CD39 and IFNγ. Samples were processed as before, recorded on a Cytek Aurora (Cytek Biosciences) and analyzed using FlowJo.

The antibody panel used for sorting of goblet cells contained antibodies against CD45, EpCAM, and CLCA1. For cell sorting, cells were stained with extracellular markers and sorted using a Cytek Aurora CS by the Flow Cytometry Core Laboratory at the Children’s Hospital of Philadelphia.

### Peptide library screening

A peptide library spanning the entire MuAstV NYU-1 capsid protein was created with 15aa peptides and 11aa overlap totaling 228 peptides. MHC I and MHC II epitopes were predicted using the Immune Epitope Database Analysis Resource (http://tools.iedb.org/main/). 50 peptides were chosen that contained the major MHC I and MHC II epitopes and synthesized by GenScript. Peptides were resuspended in DMSO to 50 mg/ml and combined for 1 mg/ml/peptide. Small intestine LPL fraction was isolated and stimulated with peptides at final concentration of 2.5 µg/ml/peptide in RPMI for 5h at 37°C. After 1h of stimulation, Protein Transport Inhibitor cocktail containing Brefeldin A and Monensin (BD) was added. Cells were analyzed by flow cytometry for IFNγ production. The MuAstV tetramer based on the final 9aa peptide was produced by the NIH Tetramer Facility.

### Single-cell RNA sequencing

About 10cm of ileum were removed, washed, digested in HBSS with 5% FCS with collagenase D (1 mg/mL) and DNase I (0.15 mg/mL, Sigma-Aldrich) for 30 min at 37°C with constant agitation and filtered through a 70μm cell strainer. Cells were counted and equal numbers from two mice were pooled. Cells were purified using Debris Removal Solution (Miltenyi Biotec) and Dead Cell Removal Kit (Miltenyi Biotec). Libraries were generated by the Single Cell Technology Core at the Children’s Hospital of Philadelphia using an 10X Genomics platform. Libraries were sequenced by the High Throughput Sequencing Core at the Children’s Hospital of Philadelphia using a NovaSeq6000 sequencing system (Illumina).

### Single-cell RNA sequencing data analysis

The Cell Ranger pipeline version 7.1. was used to demultiplex cellular barcodes and align reads against the mouse genome (mm10 ensemble). Downstream RNA-seq analysis was conducted using Seurat version 5.1. on R version 4.2.1 with the filtered RNA and HTO feature counts. Cells containing more than 10% mitochondrial DNA and fewer than 300 feature genes were filtered out during initial quality control. HTOs were normalized using centered log-ratio transformation and demultiplexed. Doublets were removed using the scDblFinder package with a calculated doublet rate of 7.5%. Regularized negative binomial regression was performed for RNA normalization using the v2 SCTransform function. Principal component analysis was conducted, and the Louvain algorithm was used for unsupervised clustering, with dimensions determined via the ElbowPlot method. UMAP representation was used to visualize the data.

Cell types were determined by a combination of unbiased clustering, canonical cell type marker signatures, and cell type annotation via the SingleR package with the ImmGenData open-source reference databases in the CellDex package. The Wilcoxon test was used to assess differentially expressed genes between clusters and treatment groups, with a Benjamini-Hochberg p-value adjustment. Differentially expressed genes were required to be expressed in a minimum of 20% of cells per group, have a log fold change of at least 1, and a p_val_adj < .01. Enrichment of Gene Ontology (GO) Biological Processes pathways was performed using the enrichGO function in the clusterProfiler package. Significant pathways required a minimum of 15 genes per pathway and p and q-values of < .01. GO categories with p < .01, q < .01, and a minimum occurrence of ≥15 genes per pathway were considered significant.

### Quantification and statistical analysis

Statistical tests were selected according to appropriate assumptions to data distribution and variance. Statistics performed as noted in figure legends and performed using R or GraphPad Prism 10 software (La Jolla, CA, USA). No statistical methods were used to predetermine sample size.

## Data availability

Sequencing data was deposited to Gene Expression Omnibus (GEO) under the accession number GSE279634.

## Acknowledgments

We wish to thank the following individuals: past and present Cadwell lab members for input and technical assistance; Dr. Matthew Weitzman, Dr. Christopher Hunter, and Dr. John Wherry for advice on the study and preparation of the manuscript; the NYU Grossman School of Medicine Flow Cytometry and Cell Sorting Core, the Flow Cytometry Core Laboratory at the Children’s Hospital of Philadelphia, the NYU Grossman School of Medicine Gnotobiotics Animal Facility, and the University of Pennsylvania Gnotobiotic Mouse Core (Penn-CHOP Microbiome Program, RRID:SCR_022384) for assistance with experiments; Blythe Philips and the University Laboratory Animal Resource (ULAR) staff at the University of Pennsylvania for veterinary advice and assistance; the Single Cell Technology Core and the High Throughput Sequencing Core at the Children’s Hospital of Philadelphia (which receive financial support from the CHOP Research Institute) for assistance with scRNA-Seq; and the NIH Tetramer Core Facility (contract number 75N93020D00005) for providing MuAstV tetramers. Figure 6G was created using BioRender. This work was supported in part by the National Institutes of Health (NIH) grants DK093668 (K.C. and S.B.K.), AI121244 (K.C.), AI140754 (K.C.), AI179896 (K.C.), DK050306 (K.C.), R01CA271245 (S.B.K.), and R44AI136141 (S.B.K.). Additional support was provided by LEO Foundation Grant LF-OC-20-000351 (S.B.K.), NYU Cancer Center Pilot grant P30CA016087 (S.B.K.), the Vilcek Fellowship (C.H.) and HHMI Gilliam Fellowship (K.Z.).

## Author contributions

C.H. and K.C. conceived the study and designed the experiments. C.H. performed, analyzed, and interpreted all the experiments. K.Z. performed the analysis of the scRNA-Seq data. E.L.A. assisted with viral detection. K.C. and S.B.K. supervised the project. C.H. and K.C. wrote the manuscript with inputs from all authors.

## References

1 Hoarau, J. J. et al. Persistent chronic inflammation and infection by Chikungunya arthritogenic alphavirus in spite of a robust host immune response. J Immunol 184, 5914–5927, doi:10.4049/jimmunol.0900255 (2010).

2 Lanford, R. E. et al. Acute hepatitis A virus infection is associated with a limited type I interferon response and persistence of intrahepatic viral RNA. Proceedings of the National Academy of Sciences of the United States of America 108, 11223–11228, doi:10.1073/pnas.1101939108 (2011).

3 Lin, W. H., Kouyos, R. D., Adams, R. J., Grenfell, B. T. & Griffin, D. E. Prolonged persistence of measles virus RNA is characteristic of primary infection dynamics. Proceedings of the National Academy of Sciences of the United States of America 109, 14989–14994, doi:10.1073/pnas.1211138109 (2012).

4 Hirsch, A. J. et al. Zika Virus infection of rhesus macaques leads to viral persistence in multiple tissues. PLoS pathogens 13, e1006219, doi:10.1371/journal.ppat.1006219 (2017).

5 Paz-Bailey, G. et al. Persistence of Zika Virus in Body Fluids - Final Report. N Engl J Med 379, 1234–1243, doi:10.1056/NEJMoa1613108 (2018).

6 Fragkoudis, R., Dixon-Ballany, C. M., Zagrajek, A. K., Kedzierski, L. & Fazakerley, J. K. Following Acute Encephalitis, Semliki Forest Virus is Undetectable in the Brain by Infectivity Assays but Functional Virus RNA Capable of Generating Infectious Virus Persists for Life. Viruses 10, doi:10.3390/v10050273 (2018).

7 Den Boon, S., et al. Ebola Virus Infection Associated with Transmission from Survivors. Emerg Infect Dis 25, 249–255, doi:10.3201/eid2502.181011 (2019).

8 Lion, T. Adenovirus persistence, reactivation, and clinical management. FEBS letters 593, 3571–3582, doi:10.1002/1873-3468.13576 (2019).

9 Owusu, D. et al. Persistent SARS-CoV-2 RNA Shedding Without Evidence of Infectiousness: A Cohort Study of Individuals With COVID-19. J Infect Dis 224, 1362–1371, doi:10.1093/infdis/jiab107 (2021).

10 Yang, B. et al. Clinical and molecular characteristics of COVID-19 patients with persistent SARS-CoV-2 infection. Nat Commun 12, 3501, doi:10.1038/s41467-021-23621-y (2021).

11 Stein, S. R. et al. SARS-CoV-2 infection and persistence in the human body and brain at autopsy. Nature 612, 758–763, doi:10.1038/s41586-022-05542-y (2022).

12 Castro, Í. A. et al. Murine parainfluenza virus persists in lung innate immune cells sustaining chronic lung pathology. Nature Microbiology, doi:10.1038/s41564-024-01805-8 (2024).

13 Ghafari, M. et al. Prevalence of persistent SARS-CoV-2 in a large community surveillance study. Nature 626, 1094–1101, doi:10.1038/s41586-024-07029-4 (2024).

14 Fischer, W. A. et al. Ebola Virus Ribonucleic Acid Detection in Semen More Than Two Years After Resolution of Acute Ebola Virus Infection. Open Forum Infect Dis 4, ofx155, doi:10.1093/ofid/ofx155 (2017).

15 Borges, V. et al. Long-Term Evolution of SARS-CoV-2 in an Immunocompromised Patient with Non-Hodgkin Lymphoma. mSphere 6, e0024421, doi:10.1128/mSphere.00244-21 (2021).

16 Chen, B., Julg, B., Mohandas, S., Bradfute, S. B. & Force, R. M. P. T. Viral persistence, reactivation, and mechanisms of long COVID. Elife 12, doi:10.7554/eLife.86015 (2023).

17 Griffin, D. E. Why does viral RNA sometimes persist after recovery from acute infections? PLoS Biol 20, e3001687, doi:10.1371/journal.pbio.3001687 (2022).

18 Tohma, K. et al. Viral intra-host evolution in immunocompetent children contributes to human norovirus diversification at the global scale. Emerg Microbes Infect 10, 1717–1730, doi:10.1080/22221751.2021.1967706 (2021).

19 Virgin, H. W., Wherry, E. J. & Ahmed, R. Redefining chronic viral infection. Cell 138, 30–50, doi:10.1016/j.cell.2009.06.036 (2009).

20 Moskophidis, D., Lechner, F., Pircher, H. & Zinkernagel, R. M. Virus persistence in acutely infected immunocompetent mice by exhaustion of antiviral cytotoxic effector T cells. Nature 362, 758–761, doi:10.1038/362758a0 (1993).

21 Zehn, D. & Wherry, E. J. Immune Memory and Exhaustion: Clinically Relevant Lessons from the LCMV Model. Adv Exp Med Biol 850, 137–152, doi:10.1007/978-3-319-15774-0_10 (2015).

22 Sharpe, A. H., Wherry, E. J., Ahmed, R. & Freeman, G. J. The function of programmed cell death 1 and its ligands in regulating autoimmunity and infection. Nat Immunol 8, 239–245, doi:10.1038/ni1443 (2007).

23 Nakamoto, N. et al. Functional restoration of HCV-specific CD8 T cells by PD-1 blockade is defined by PD-1 expression and compartmentalization. Gastroenterology 134, 1927–1937, 1937 e1921-1922, doi:10.1053/j.gastro.2008.02.033 (2008).

24 Tomov, V. T. et al. Differentiation and Protective Capacity of Virus-Specific CD8(+) T Cells Suggest Murine Norovirus Persistence in an Immune-Privileged Enteric Niche. Immunity 47, 723–738 e725, doi:10.1016/j.immuni.2017.09.017 (2017).

25 Tomov, V. T. et al. Persistent enteric murine norovirus infection is associated with functionally suboptimal virus-specific CD8 T cell responses. J Virol 87, 7015–7031, doi:10.1128/JVI.03389-12 (2013).

26 Strine, M. S. et al. Intestinal tuft cell immune privilege enables norovirus persistence. Sci Immunol 9, eadi7038, doi:10.1126/sciimmunol.adi7038 (2024).

27 Barber, D. L. et al. Restoring function in exhausted CD8 T cells during chronic viral infection. Nature 439, 682–687, doi:10.1038/nature04444 (2006).

28 Nice, T. J. et al. Interferon-lambda cures persistent murine norovirus infection in the absence of adaptive immunity. Science 347, 269–273, doi:10.1126/science.1258100 (2015).

29 Knipe, D. M. & Howley, P. M. Fields virology. 6th edn, (Wolters Kluwer/Lippincott Williams & Wilkins Health, 2013).

30 Koopmans, M. P., Bijen, M. H., Monroe, S. S. & Vinje, J. Age-stratified seroprevalence of neutralizing antibodies to astrovirus types 1 to 7 in humans in The Netherlands. Clin Diagn Lab Immunol 5, 33–37, doi:10.1128/CDLI.5.1.33-37.1998 (1998).

31 Farkas, T., Fey, B., Keller, G., Martella, V. & Egyed, L. Molecular detection of novel astroviruses in wild and laboratory mice. Virus Genes 45, 518–525, doi:10.1007/s11262-012-0803-0 (2012).

32 Yokoyama, C. C. et al. Adaptive immunity restricts replication of novel murine astroviruses. J Virol 86, 12262–12270, doi:10.1128/JVI.02018-12 (2012).

33 Ingle, H. et al. Murine astrovirus tropism for goblet cells and enterocytes facilitates an IFN-lambda response in vivo and in enteroid cultures. Mucosal Immunol, doi:10.1038/s41385-021-00387-6 (2021).

34 Cortez, V. et al. Astrovirus infects actively secreting goblet cells and alters the gut mucus barrier. Nat Commun 11, 2097, doi:10.1038/s41467-020-15999-y (2020).

35 Cortez, V. et al. Characterizing a Murine Model for Astrovirus Using Viral Isolates from Persistently Infected Immunocompromised Mice. J Virol 93, doi:10.1128/JVI.00223-19 (2019).

36 Marvin, S. A. et al. Type I Interferon Response Limits Astrovirus Replication and Protects against Increased Barrier Permeability In Vitro and In Vivo. Journal of virology 90, 1988–1996, doi:10.1128/JVI.02367-15 (2016).

37 Pernold, K. et al. Towards large scale automated cage monitoring - Diurnal rhythm and impact of interventions on in-cage activity of C57BL/6J mice recorded 24/7 with a non-disrupting capacitive-based technique. PLoS One 14, e0211063, doi:10.1371/journal.pone.0211063 (2019).

38 Jumper, J. et al. Highly accurate protein structure prediction with AlphaFold. Nature 596, 583–589, doi:10.1038/s41586-021-03819-2 (2021).

39 Lanning, S. et al. Structure and immunogenicity of the murine astrovirus capsid spike. The Journal of general virology 104, doi:10.1099/jgv.0.001913 (2023).

40 Sanchez-Fauquier, A. et al. Characterization of a human astrovirus serotype 2 structural protein (VP26) that contains an epitope involved in virus neutralization. Virology 201, 312–320, doi:10.1006/viro.1994.1296 (1994).

41 Bass, D. M. & Upadhyayula, U. Characterization of human serotype 1 astrovirus-neutralizing epitopes. J Virol 71, 8666–8671, doi:10.1128/JVI.71.11.8666-8671.1997 (1997).

42 Dallari, S. et al. Enteric viruses evoke broad host immune responses resembling those elicited by the bacterial microbiome. Cell host & microbe, doi:10.1016/j.chom.2021.03.015 (2021).

43 Jiang, Y., Li, Y. & Zhu, B. T-cell exhaustion in the tumor microenvironment. Cell death & disease 6, e1792, doi:10.1038/cddis.2015.162 (2015).

44 Karginov, T. A., Menoret, A. & Vella, A. T. Optimal CD8(+) T cell effector function requires costimulation-induced RNA-binding proteins that reprogram the transcript isoform landscape. Nat Commun 13, 3540, doi:10.1038/s41467-022-31228-0 (2022).

45 Martin-Cofreces, N. B., Baixauli, F. & Sanchez-Madrid, F. Immune synapse: conductor of orchestrated organelle movement. Trends Cell Biol 24, 61–72, doi:10.1016/j.tcb.2013.09.005 (2014).

46 Hu, M. et al. DDX5: an expectable treater for viral infection-a literature review. Ann Transl Med 10, 712, doi:10.21037/atm-22-2375 (2022).

47 Cadwell, K. et al. Virus-plus-susceptibility gene interaction determines Crohn’s disease gene Atg16L1 phenotypes in intestine. Cell 141, 1135–1145, doi:10.1016/j.cell.2010.05.009 (2010).

48 Subramani, S. & Malhotra, V. Non-autophagic roles of autophagy-related proteins. EMBO Rep 14, 143–151, doi:10.1038/embor.2012.220 (2013).

49 Hwang, S. et al. Nondegradative role of Atg5-Atg12/ Atg16L1 autophagy protein complex in antiviral activity of interferon gamma. Cell host & microbe 11, 397–409, doi:10.1016/j.chom.2012.03.002 (2012).

50 Hardy, R. S., Raza, K. & Cooper, M. S. Therapeutic glucocorticoids: mechanisms of actions in rheumatic diseases. Nat Rev Rheumatol 16, 133–144, doi:10.1038/s41584-020-0371-y (2020).

51 Phan, T. G. et al. The fecal viral flora of wild rodents. PLoS pathogens 7, e1002218, doi:10.1371/journal.ppat.1002218 (2011).

52 Rosshart, S. P. et al. Laboratory mice born to wild mice have natural microbiota and model human immune responses. Science 365, doi:10.1126/science.aaw4361 (2019).

53 Fay, E. J. et al. Natural rodent model of viral transmission reveals biological features of virus population dynamics. J Exp Med 219, doi:10.1084/jem.20211220 (2022).

54 Cruz-Topete, D. & Cidlowski, J. A. One hormone, two actions: anti- and pro-inflammatory effects of glucocorticoids. Neuroimmunomodulation 22, 20–32, doi:10.1159/000362724 (2015).

55 Hong, J. Y. et al. Long-Term Programming of CD8 T Cell Immunity by Perinatal Exposure to Glucocorticoids. Cell 180, 847–861 e815, doi:10.1016/j.cell.2020.02.018 (2020).

56 Tehseen, A. et al. Glucocorticoid-mediated Suppression of Effector Programming Assists the Memory Transition of Virus-specific CD8+ T Cells. J Immunol, doi:10.4049/jimmunol.2300513 (2024).

57 Jamieson, A. M., Yu, S., Annicelli, C. H. & Medzhitov, R. Influenza virus-induced glucocorticoids compromise innate host defense against a secondary bacterial infection. Cell host & microbe 7, 103–114, doi:10.1016/j.chom.2010.01.010 (2010).

58 Schneider, K. M. et al. The enteric nervous system relays psychological stress to intestinal inflammation. Cell 186, 2823–2838 e2820, doi:10.1016/j.cell.2023.05.001 (2023).

59 Group, R. C. et al. Dexamethasone in Hospitalized Patients with Covid-19. N Engl J Med 384, 693–704, doi:10.1056/NEJMoa2021436 (2021).

60 Vashist, S., Urena, L. & Goodfellow, I. Development of a strand specific real-time RT-qPCR assay for the detection and quantitation of murine norovirus RNA. J Virol Methods 184, 69–76, doi:10.1016/j.jviromet.2012.05.012 (2012).

61 Dong, J., Dong, L., Mendez, E. & Tao, Y. Crystal structure of the human astrovirus capsid spike. Proceedings of the National Academy of Sciences of the United States of America 108, 12681–12686, doi:10.1073/pnas.1104834108 (2011).

